# A two-membrane gateway for monocarboxylates couples host glycolysis to *Toxoplasma* mitochondrial fitness and virulence

**DOI:** 10.64898/2026.02.23.707530

**Authors:** Melanie Key, Steven Joseph, Zhicheng Dou

**Author notes:** Corresponding author: Zhicheng Dou.

## Abstract

Intracellular pathogens must coordinate their metabolism with nutrient supplies from the host cell, yet the specific metabolites and transport pathways that sustain parasite bioenergetics remain incompletely defined. In the apicomplexan parasite *Toxoplasma gondii*, infection increases host glycolytic flux and elevates cytosolic lactate and pyruvate, suggesting that these intermediates are co-opted as carbon and energy sources. Here, we show that *T. gondii* imports host-derived lactate and pyruvate across both the parasitophorous vacuole membrane and the parasite plasma membrane to maintain mitochondrial function, extracellular survival, and acute virulence. Using a hexokinase knockout (Δ*hk*) to abolish endogenous pyruvate production, we find that parasites preserve basal oxygen consumption but become strictly dependent on exogenous monocarboxylates to stimulate mitochondrial respiration. By disrupting the parasite formate-nitrite transporters TgFNT1-3, we identify TgFNT1 and TgFNT2 as the principal monocarboxylate transporters required for lactate- and pyruvate-driven respiratory responses. Furthermore, genetic ablation of TgGRA17, a parasitophorous vacuole pore protein, compromises the growth advantage conferred by elevated exogenous lactate, implicating this pore as the entry route for host-derived monocarboxylates into the vacuole. Conversely, host cells lacking the monocarboxylate exporter MCT1 accumulate cytosolic lactate/pyruvate and enhance parasite growth, linking host monocarboxylate export to parasite fitness. When both endogenous pyruvate production and exogenous uptake are disrupted, parasites display severely reduced mitochondrial basal respiratory capacity, membrane potential, ATP levels, extracellular survival, and virulence in mice. Collectively, these findings define a dual-step pyruvate acquisition pathway in *T. gondii* and reveal host monocarboxylates as critical fuels that buffer parasite bioenergetic stress during infection.

**Significance Statement:** Intracellular parasites rely on host nutrients to power their metabolism, yet the routes by which these metabolites cross the membranes between host cytosol and parasite mitochondria are not well defined. Here, we show that *Toxoplasma gondii* exploits host glycolysis by importing lactate and pyruvate to sustain mitochondrial function and virulence. We identify a two-step pathway in which these monocarboxylates cross the parasitophorous vacuole via the pore GRA17 and then enter the parasite through the formate-nitrite transporters TgFNT1/2. Blocking both endogenous glycolysis and this exogenous pyruvate supply disables parasite mitochondrial fitness, extracellular survival, and virulence. These findings reveal a fundamental strategy of metabolic plasticity in apicomplexan parasites using a multi-membrane nutrient gateway that couples host glycolysis to parasite bioenergetics.

## Introduction

Apicomplexan parasites such as *Toxoplasma gondii* and *Plasmodium falciparum* replicate within host cells inside a specialized compartment, namely the parasitophorous vacuole (PV). The PV is derived from invaginated host plasma membrane and extensively remodeled by parasite-secreted proteins, creating a niche that physically separates parasites from the host cytosol. Within this compartment, parasites must secure a continuous influx of nutrients to support growth, mitochondrial function, and the production of virulence factors. How intracellular parasites integrate nutrients produced by the host with their own metabolic pathways remains a central question in infection biology.

*Toxoplasma gondii* profoundly rewires host cell metabolism during infection. In particular, upregulation of host hexokinase II increases glycolytic flux and elevates cytosolic concentrations of lactate and pyruvate (1, 2). These end products of glycolysis are abundant, diffusible metabolites with the potential to serve as carbon and energy sources for intracellular parasites. Indeed, exogenous lactate can rescue growth of *T. gondii* mutants lacking pyruvate kinase (TgPYK1) (3), which also serves as a key enzyme in the production of endogenous pyruvate pool, suggesting that the parasite can acquire and metabolize extracellular glycolytic intermediates. However, the specific routes by which lactate and pyruvate traverse both the parasitophorous vacuole membrane (PVM) and the parasite plasma membrane, and the extent to which these host-derived monocarboxylates contribute to parasite bioenergetics and virulence, are not well defined.

To overcome the diffusion barrier imposed by the PVM, *T. gondii* secretes dense granule proteins that assemble into nutrient-permeable pores. Among these, TgGRA17 and TgGRA23 form channels that increase PVM conductance and allow small solutes (<1.3 kDa) to cross between the host cytosol and PV lumen. Disruption of TgGRA17 reduces PVM permeability, causes swollen vacuoles, and diminishes acute virulence in mice, implicating this protein as a major conduit for nutrient exchange (4, 5). In addition, TgGRA23 was found to compensate for the loss of TgGRA17, since the simultaneous deletion of both genes is lethal in *Toxoplasma*. Two recent studies identified two additional dense granule proteins, TgGRA47 and TgGRA72, that are involved in pore formation on the PVM, thereby affecting the PVM permeability (6, 7). In particular, the deletion of TgGRA72 in Δ*gra17* resulted in synthetic lethality, suggesting that TgGRA72 can compensate for the loss of TgGRA17, underscoring the importance of the small molecule-accessible pore system for tachyzoite survival.

Once small solutes reach the PV lumen, they must still cross the parasite plasma membrane before utilization. *T. gondii* encodes multiple transporters for sugars (*e.g.,* TgGT1 and TgST2 (8)) and amino acids (*e.g.,* TgNPT1 (9) and TgApiAT family (10)), as well as a family of formate-nitrite transporter (FNT)-like proteins, TgFNT1-3 (11–13). Based on the mass spectrometry datasets in ToxoDB, TgFNT1 (TGGT1_209800) and TgFNT2 (TGGT1_292110) are detected in tachyzoites, whereas TgFNT3 (TGGT1_229170) expression is under detection limits but was seen in bradyzoites (12). In addition to formate, TgFNT1-3 transporters have been reported to transport lactate across the plasma membrane when expressed in yeast (11–13). A previous work showed that purified extracellular *Toxoplasma* parasites can incorporate the lactate and acidify the cytoplasm (11). In bacteria, the FNT ortholog, namely FocA protein, can operate bidirectionally (14, 15). A recent work documented that the intracellular parasites can export lactate and pyruvate from their cytoplasm to PV lumen for acidification via TgFNT1 and TgFNT2, but deletion of these transporters produces only modest defects in growth and egress (13). These observations raise the possibility that TgFNTs might also mediate uptake of lactate or pyruvate from the PV lumen in addition to acting as exporters.

*Toxoplasma gondii* possesses a complete glycolytic pathway to generate an endogenous pool of glycolytic metabolites (16, 17). Hexokinase (TgHK, TGGT1_265450) serves as the first enzyme within this glucose catabolism and is dispensable in *T. gondii* (18). The mutant lacking TgHK (Δ*hk*) does not exhibit severe growth defects under standard, glucose-rich culture conditions, likely because it engages alternative pathways, including glutamine anaplerosis via the GABA shunt (8), to support the tricarboxylic acid (TCA) cycle. However, the Δ*hk* mutant forms significantly fewer cysts than wildtype, yet it still replicates in glutamine-depleted medium (18). These observations highlight the metabolic plasticity of *T. gondii* and raise the possibility that parasites can supplement or bypass their own glycolysis by importing other host-derived carbon sources, particularly under conditions where glucose and glutamine availability or glycolytic flux are restricted.

Taken together, existing work suggests a model in which *T. gondii* could draw on two distinct sources of pyruvate: (i) pyruvate generated endogenously by parasite glycolysis, and (ii) pyruvate derived from host glycolysis, accessed as lactate and pyruvate through PVM pores and parasite plasma membrane transporters. This model raises two key questions. First, do tachyzoites directly import host-derived lactate and pyruvate across both the PVM and parasite plasma membrane, and if so, which proteins mediate these fluxes? Second, how important are these exogenous monocarboxylates, relative to endogenously produced pyruvate, for mitochondrial function, extracellular survival, and virulence during infection?

Here, we use molecular genetics, bioenergetic profiling, and *in vivo* infection models to dissect the contribution of host-derived lactate and pyruvate to *T. gondii* pathogenesis. Using a TgHK knockout (Δ*hk*) to disable endogenous glycolytic pyruvate production, we show that parasites maintain basal mitochondrial respiration but become highly responsive to exogenous lactate and pyruvate. We utilize this sensitized background to define the roles of TgFNT1-3 and TgGRA17 in monocarboxylate transport. Genetic ablation of TgFNT1 and TgFNT2 abolishes lactate- and pyruvate-driven stimulation of mitochondrial respiration, while deletion of TgGRA17 reduces the growth advantage conferred by elevated host lactate. Host cells lacking the monocarboxylate exporter MCT1 accumulate cytosolic lactate/pyruvate and support enhanced parasite growth, linking host monocarboxylate export to parasite fitness. Finally, by combining Δ*hk* with TgFNT and TgGRA17 deletions, we uncouple endogenous and exogenous pyruvate sources and show that simultaneous disruption of both pathways severely impairs mitochondrial basal respiratory capacity, membrane potential, ATP production, extracellular survival, and acute virulence in mice. These findings define a dual-source pyruvate acquisition system in *T. gondii* and identify host-derived glycolytic intermediates as critical fuels that buffer parasite bioenergetic stress during infection.

## Results

### 1. *Toxoplasma* actively incorporates and metabolizes extracellular glycolytic metabolites

Prior studies show that *Toxoplasma* infection elevates host glycolysis (1, 2) and that a pyruvate kinase-deficient parasite can recover growth when the medium is supplemented with lactate (3). These observations suggest that parasites can access extracellular glycolytic end products, but they did not demonstrate whether parasites directly metabolize these metabolites.

Because pyruvate links glycolysis to mitochondrial oxidative phosphorylation, we sought to distinguish lactate produced endogenously by parasites from lactate acquired from the extracellular environment. We therefore deleted parasite hexokinase (TgHK) to generate a Δ*hk* mutant. To confirm that loss of TgHK abolishes parasite-derived lactate secretion, extracellular parasites were incubated in DMEM supplemented with 10 mM glucose for 1 h, and lactate in the supernatant was quantified by a fluorometric assay. Compared with WT parasites, Δ*hk* secreted negligible lactate (**Fig. 1A**). Consistently, treating WT parasites with 5 mM 2-deoxy-D-glucose (2-DG), a hexokinase inhibitor, similarly suppressed lactate production (**Fig. 1A**). Thus, Δ*hk* provides a system to evaluate the contribution of imported lactate independently of parasite-derived lactate.

**Figure 1.**
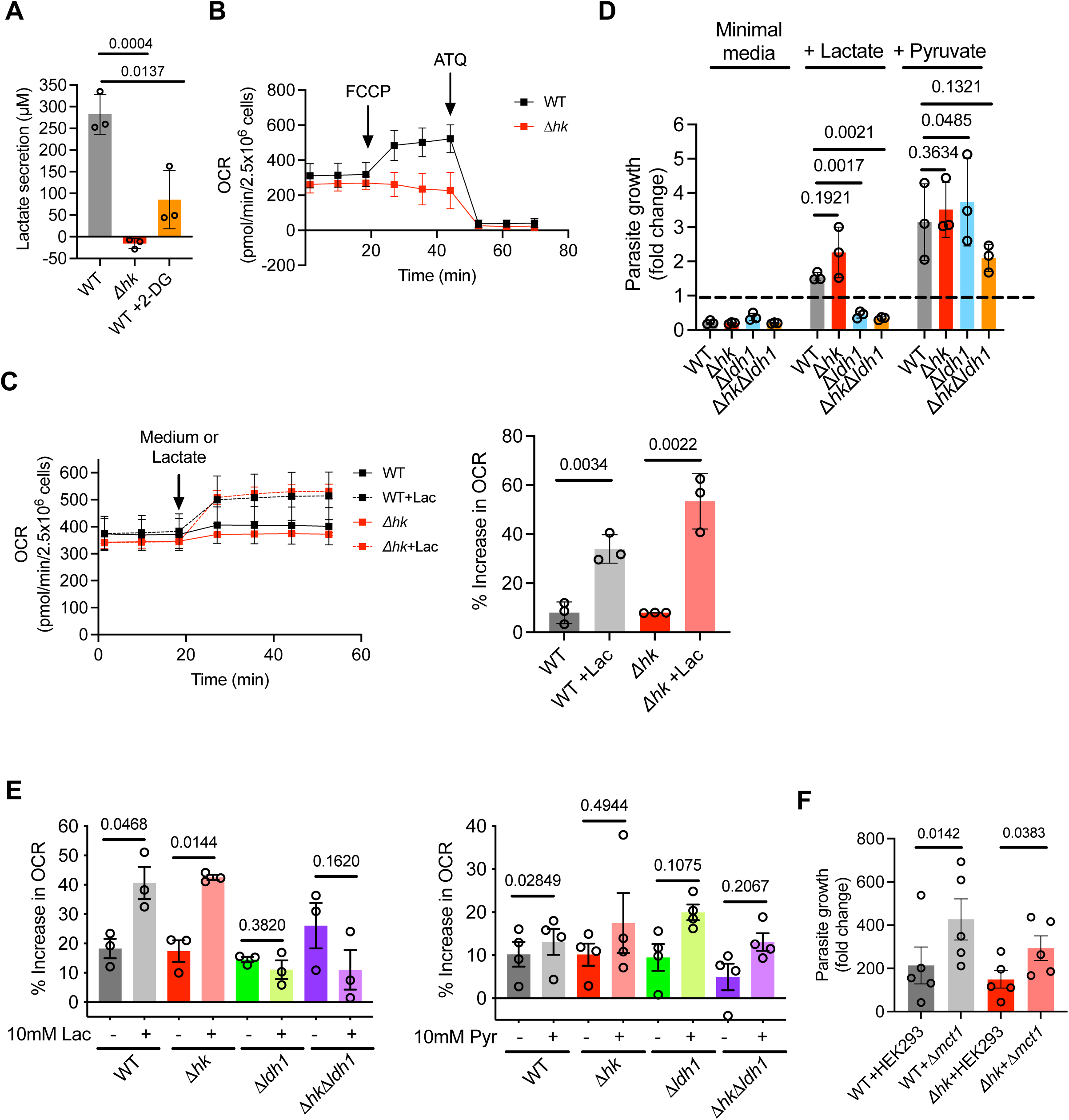
Parasites lacking hexokinase show increased dependence on exogenous lactate. (A) Lactate secretion quantified by fluorometric assay. Purified extracellular tachyzoites were incubated for 1 h in DMEM containing 10 mM glucose prior to measurement of released lactate. WT parasites were treated with 5 mM 2-deoxy-D-glucose (2-DG) before lactate quantification. (B) Representative Seahorse oxygen consumption rate (OCR) graph for WT and Δ*hk* parasites. FCCP was injected to uncouple mitochondrial membrane potential and drive maximal respiration, followed by atovaquone (ATQ) to inhibit complex III. (C) Acute lactate stimulation of mitochondrial respiration. Representative OCR traces following injection of base medium alone or base medium supplemented with 10 mM lactate. Lactate-stimulated OCR responses are shown as percent increase over baseline from 3 biological replicates. (D) Parasite growth under defined carbon sources measured by luciferase assay. WT, Δ*hk*, Δ*ldh1*, and Δ*hk*Δ*ldh1* parasites were grown in HFFs for 48 h in carbon source-free minimal medium or the same medium supplemented with 10 mM lactate or 10 mM pyruvate. (E) Acute OCR responses of strains listed in Panel D following injection of base medium alone or base medium supplemented with 10 mM lactate or 10 mM pyruvate, plotted as percent increase over baseline OCR. (F) Replication of WT parasites in parental HEK293 cells versus HsMCT1-deficient (Δ*mct1*) HEK293 cells, measured 96 h post-infection by luciferase assay. Data are presented as mean ± SD or SEM from 3-5 independent biological replicates as indicated in each figure. Statistical significance was determined by two-tailed unpaired Student’s *t*-test (Panel A and C) and two-tailed paired Student’s *t*-test (Panel D, E, and F).

Since pyruvate enters the mitochondria for metabolism, we measured the impact of endogenously produced lactate on mitochondrial respiration rates by quantifying oxygen consumption rate (OCR) in WT and Δ*hk* parasites using an Agilent Seahorse assay. Our data showed that the basal OCR was comparable between WT and Δ*hk* parasites (**Fig. 1B**), indicating that mitochondrial respiration is largely maintained despite impaired glycolytic flux. To test whether parasites directly metabolize extracellular monocarboxylates, we acutely injected 10 mM lactate into Seahorse assay medium and monitored OCR. WT parasites showed a rapid increase in OCR upon lactate addition (**Fig. 1C**), consistent with immediate mitochondrial utilization of exogenous lactate. Notably, Δ*hk* parasites exhibited an even larger OCR increase than WT after lactate addition (**Fig. 1C**), suggesting enhanced dependence on extracellular substrates when endogenous pyruvate is limiting. Acute pyruvate addition produced a similar, though weaker, OCR increase (**Supplemental Fig. 1A and B**).

### 2. Exogenous glycolytic metabolites promote intracellular parasite growth

Lactate supplementation can rescue growth of a pyruvate kinase-deficient mutant (3), but it remains unclear whether parasites directly use exogenous lactate or its derived metabolites. Since lactate must be converted to pyruvate prior to entry into central carbon metabolism, we tested whether lactate utilization requires the parasite’s lactate dehydrogenase (LDH). *Toxoplasma* encodes two LDHs (19), but only TgLDH1 (TGGT1_232350) is predominantly expressed during acute infection (20). Therefore, we deleted *TgLDH1* in WT::*NLuc* and *Δhk*::*NLuc* backgrounds to generate Δ*ldh1*::*NLuc* and Δ*hk*Δ*ldh1*::*NLuc* (**Supplemental Fig. 2A-C**). The resulting *TgLDH1* null mutant showed a ∼95% reduction in lactate secretion (**Supplemental Fig. 3**), indicating that TgLDH1 is the primary enzyme converting pyruvate to lactate in tachyzoites and suggesting that Δ*ldh1* parasites cannot effectively utilize imported exogenous lactate.

To further test this speculation, WT, TgHK- and/or TgLDH1-deletion parasites were grown in minimal DMEM lacking carbon sources or supplemented with 10 mM lactate or pyruvate. At 48 h post-infection, all strains grew in pyruvate-supplemented medium; in contrast, strains lacking *TgLDH1* failed to grow in lactate-supplemented medium (**Fig. 1D**). Consistently, Seahorse analysis showed that Δ*ldh1* and Δ*hk*Δ*ldh1* increased OCR in response to pyruvate, but not lactate (**Fig. 1E**), supporting the conclusion that imported lactate must be converted to pyruvate via TgLDH1 prior to utilization.

We next asked whether increasing cytosolic monocarboxylates in host cells enhances parasite growth. Lactate/pyruvate in the medium must cross the host plasma membrane and the parasitophorous vacuole membrane before reaching the parasite. To elevate host cytosolic monocarboxylates, we deleted the mammalian monocarboxylate transporter MCT1 in HEK293 cells to generate Δ*mct1* cells. MCT1 contributes to lactate/pyruvate export (21), therefore, its disruption is expected to increase intracellular lactate/pyruvate. CRISPR-mediated deletion of both MCT1 alleles abolished MCT1 protein expression by immunoblot (**Supplemental Fig. 4A**) and reduced medium acidification relative to WT HEK293 cells (**Supplemental Fig. 4B**). Using the FRET-based lactate biosensor named Laconic (22), Δ*mct1* cells exhibited an ∼1.5-fold higher cytosolic lactate signal than WT HEK293 cells (**Supplemental Fig. 4C**), corresponding to an approximate 10-fold increase in intracellular lactate based on a standard curve reported previously in the literature (22). When WT::*NLuc* or Δ*hk*::*NLuc* parasites were grown in Δ*mct1* cells, intracellular growth increased by ∼2-fold compared with growth in WT HEK293 cells (**Fig. 1F**), indicating that parasites can access host cytosolic lactate to support replication.

### 3. TgFNT1 and TgFNT2 mediate import of exogenous glycolytic metabolites across the parasite plasma membrane as fuels for mitochondrial metabolism

*T. gondii* encodes three plasma-membrane formate/nitrite transporter (FNT)-like proteins reported to transport monocarboxylates bidirectionally (*e.g.,* lactate, formate, nitrite) (11–13). TgFNT1 and TgFNT2 are expressed during both major stages, whereas TgFNT3 has been proposed to be bradyzoite-enriched based on transcriptomic data (11, 12). Previous work showed that loss of TgFNT1 reduced cytoplasmic acidification when Δ*fnt1* parasites were incubated in 10 mM lactate, but this observation provides only indirect evidence to suggest that TgFNT1 is a major lactate importer (11). Therefore, these data do not clearly establish whether TgFNT1 is the sole transporter responsible for importing glycolytic intermediates, nor do they define the metabolic fate of these molecules after entry into the parasite cytoplasm. We hypothesized that one or more TgFNTs mediate lactate/pyruvate import into parasites.

We generated single knockouts (Δ*fnt1*, Δ*fnt2*, Δ*fnt3*) in the Δ*hk* parasites to generate three individual cell lines, Δ*hk*Δ*fnt1*, Δ*hk*Δ*fnt2,* and Δ*hk*Δ*fnt3* (**Supplemental Fig. 5A and Supplemental Fig. 6A-B**), and quantified lactate- or pyruvate-stimulated OCR increases. The deletion of TgHK is expected to help eliminate the background OCR from endogenously produced pyruvate. Loss of any single TgFNT did not eliminate the OCR response to exogenous lactate or pyruvate (**Supplemental Fig. 7A**), suggesting redundant functions of these TgFNT homologs. We therefore deleted TgFNT1 and TgFNT2 together to generate Δ*hk*Δ*fnt1/2* (**Supplemental Fig. 5B**). Although TgFNT3 is not detectable in tachyzoites under standard conditions (12), loss of TgFNT1/2 could potentially induce compensatory upregulation of TgFNT3. Therefore, we additionally generated Δ*hk*Δ*fnt1/2/3* triple-knockout parasites to ablate TgFNT expression and eliminate TgFNT-mediated uptake of exogenous carboxylates (**Supplemental Fig. 5C**). Strikingly, deletion of TgFNT1/2 completely abolished the OCR increase upon lactate addition (**Fig. 2A**) and similarly reduced the OCR response to pyruvate (**Fig. 2A**). To validate these observations, we created the Δ*fnt1*, Δ*fnt1/2*, and Δ*fnt1/2/3* parasites (**Supplemental Fig. 8A-C**) and repeated the OCR determination with addition of lactate and pyruvate. We observed a similar trend: loss of TgFNT1/2 completely abolishes the OCR increase, albeit to a lesser extent (**Supplemental Fig. 7B**). Overall, these results identify TgFNT1 and TgFNT2 as the principal routes for importing exogenous lactate/pyruvate across the tachyzoite plasma membrane before they are utilized in mitochondria for further metabolism.

**Figure 2.**
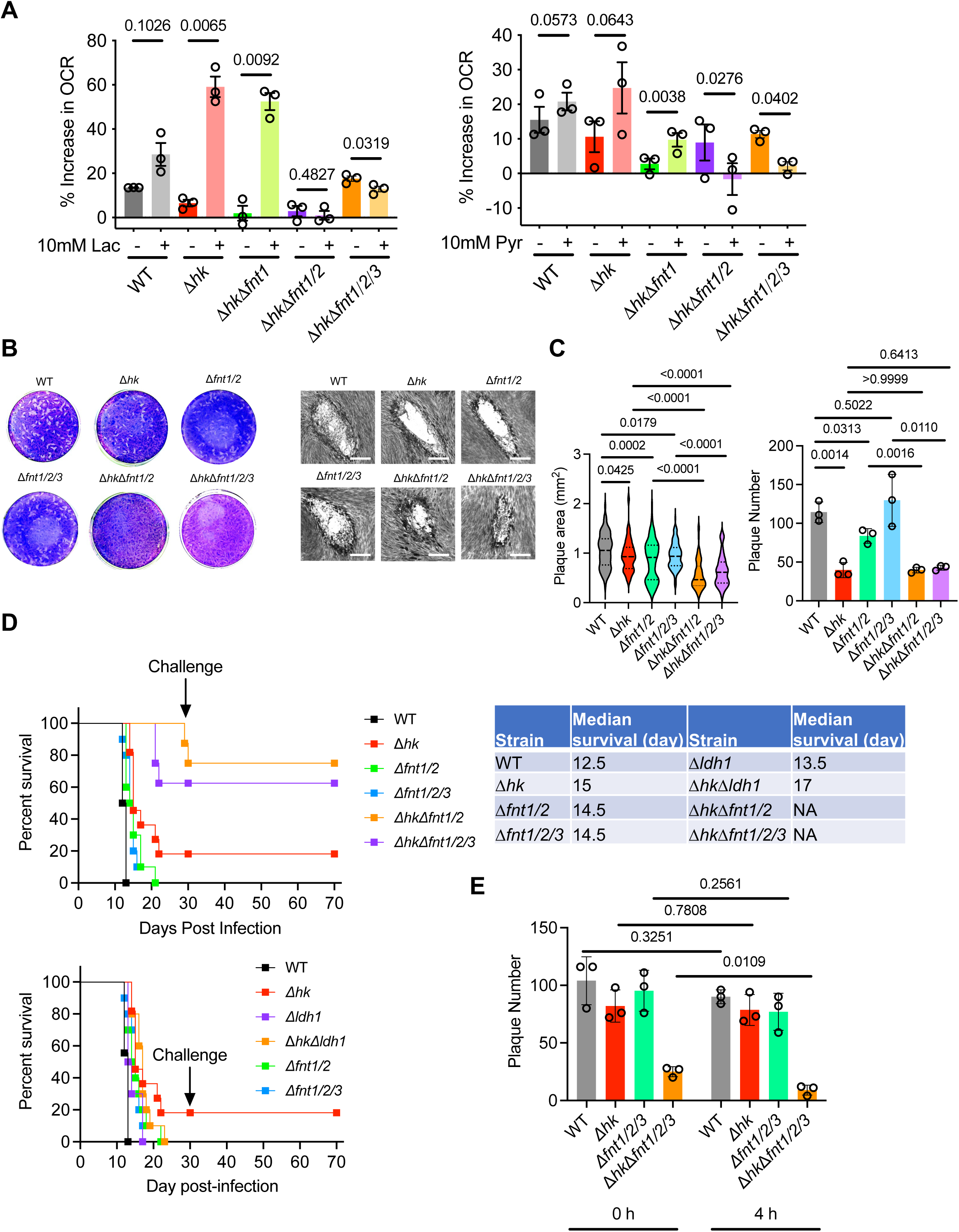
TgFNT1 and TgFNT2 mediate glycolytic metabolite uptake and contribute to acute virulence. (A) Lactate- or pyruvate-stimulated OCR in extracellular parasites lacking TgFNT transporter(s) measured by Seahorse assay. Parasites were adhered with Cell-Tak and assayed in base medium containing 10 mM glucose and 4 mM glutamine, followed by acute injection of 10 mM lactate or 10 mM pyruvate. Responses are plotted as percent increase relative to basal OCR. (B) Representative plaque assay images from HFF monolayers infected with ∼200 tachyzoites per well and cultured for 7 d prior to crystal violet staining. Bar = 500 µm. (C) Quantification of plaque area (≥20 plaques per replicate) and plaque number from three independent biological replicates. (D) Survival of CD-1 mice (n = 10 per strain; 5 male and 5 female) inoculated subcutaneously with 100 tachyzoites of the indicated strains and monitored for 30 d. Surviving mice were assessed for seroconversion and rechallenged with 1,000 WT tachyzoites to evaluate acquired immunity. Survival curves were compared using the log-rank (Mantel-Cox) test. (E) Extracellular survival assessed by plaque formation after parasites were incubated in 1× PBS for 0 h or 4 h prior to plating on HFF monolayers. Plaques were quantified for comparison. Data are presented as mean ± SD or SEM with 3 independent biological replicates denoted in each panel. Unless otherwise noted, statistical significance was determined by two-tailed unpaired Student’s *t*-test (Panel C and E) and two-tailed paired Student’s *t*-test (Panel A).

### 4. Import of exogenous glycolytic metabolites is critical for parasite fitness when endogenous production is impaired

Δ*hk* parasites cannot produce pyruvate/lactate efficiently from glucose, whereas Δ*fnt1/2* (or Δ*fnt1/2/3*) parasites are defective in importing exogenous lactate/pyruvate. Thus, Δ*hk*Δ*fnt1/2* (or Δ*hk*Δ*fnt1/2/3*) parasites lack both endogenous production and exogenous uptake, enabling us to test the contribution of these pathways to parasite fitness.

In plaque assays, Δ*hk* parasites formed moderately smaller plaques than WT, whereas Δ*fnt1/2* (or Δ*fnt1/2/3*) parasites produced plaques comparable to WT (**Fig. 2B-C**). Deletion of individual TgFNT genes also did not reduce plaque size relative to the corresponding parental strains (**Supplemental Fig. 9A-B**). In contrast, Δ*hk*Δ*fnt1/2* and Δ*hk*Δ*fnt1/2/3* parasites exhibited ∼50% smaller plaques and produced fewer plaques overall (**Fig. 2B-C**). To define which step(s) of the lytic cycle were affected, we assessed invasion, replication, and egress. The Δ*hk* mutant showed a significant invasion defect by ∼50% reduction, whereas Δ*fnt1/2/3* parasites displayed only a modest decrease by ∼20% compared with WT. Disrupting both pyruvate-producing routes further decreased invasion efficiency (**Supplemental Fig. 9C**). By contrast, intracellular replication and egress were not detectably altered in these mutants (**Supplemental Fig. 9D-E**). Together, these phenotypes indicate a strong synthetic fitness defect when both acquisition pathways are disabled.

To assess acute virulence, 10 CD-1 mice (5 males and 5 females) were inoculated subcutaneously with 100 tachyzoites per strain and monitored for survival for 30 days. Mice infected with Δ*hk*, Δ*fnt1/2*, or Δ*fnt1/2/3* parasites exhibited delayed mortality relative to WT; notably, one Δ*hk*-infected mouse survived the acute phase (**Fig. 2D**). In contrast, Δ*hk*Δ*fnt1/2* and Δ*hk*Δ*fnt1/2/3* parasites exhibited remarkably attenuated virulence, and all infected mice survived through the assessment period. However, in accordance with our IACUC protocol, mice that lost >20% of their starting body weight were euthanized, resulting in apparent survival rates of ∼80% and ∼70% for mice infected with Δ*hk*Δ*fnt1/2* and Δ*hk*Δ*fnt1/2/3*, respectively (**Fig. 2D**). Since TgFNT deletions block uptake of both lactate and pyruvate, whereas TgLDH1 deletion specifically blocks lactate utilization, we compared acute virulence of Δ*ldh1* and Δ*hk*Δ*ldh1* parasites. Δ*ldh1* parasites displayed slightly delayed mortality compared with WT, and Δ*hk*Δ*ldh1* parasites were further attenuated relative to Δ*ldh1* (**Fig. 2D**). The animal data suggest that uptake of exogenous glycolytic intermediates contributes to the parasite’s acute virulence. Notably, selectively blocking lactate utilization in Δ*ldh1*, rather than pyruvate utilization, resulted in a less pronounced loss of acute virulence than that observed for Δ*fnt1/2* or Δ*fnt1/2/3* mutants that halt import of both lactate and pyruvate (**Fig. 2D**), indicating that both metabolites are likely imported and contribute to parasite virulence.

Finally, because exogenous lactate/pyruvate can be metabolized by parasites, we asked whether these substrates support extracellular survival. Freshly egressed parasites were filter-purified and incubated in complete D10% medium for 4 h prior to plating for plaque formation, while parasites plated immediately after purification served as controls. WT, Δ*hk*, and Δ*fnt1/2/3* parasites retained plaque-forming efficiency after 4 h, whereas Δ*hk*Δ*fnt1/2/3* parasites lacking both endogenous and exogenous pyruvate/lactate acquisition showed reduced survival (**Fig. 2E**). Together, these results indicate that uptake of exogenous glycolytic end products provides an important fitness advantage, particularly when endogenous glycolysis-derived pyruvate is limiting.

### 5. TgGRA17 contributes to acquisition of host-derived glycolytic metabolites across the parasitophorous vacuole membrane

Before reaching the parasite cytosol, host-derived monocarboxylates must traverse the parasitophorous vacuole membrane (PVM). Several dense granule proteins (*e.g.,* TgGRA17, TgGRA23, TgGRA47, TgGRA72) have been implicated in forming pores that permit diffusion of small solutes (<1.3 kDa) across the PVM (4–7, 23). TgGRA17 appears to be a major component of this permeability pathway, and its deletion results in swollen vacuoles (4).

Because Δ*hk* parasites rely more heavily on extracellular/host-derived substrates, we hypothesized that deleting TgGRA17 in the Δ*hk* background would restrict nutrient passage across the PVM, including limited access to exogenous glycolytic metabolites and further impairing parasite fitness. Using CRISPR-based genome editing, we generated a Δ*hk*Δ*gra17* strain (**Supplemental Fig. 10**). Loss of *TgGRA17* in the Δ*hk* background resulted in fewer and smaller plaques compared with loss of *TgGRA17* in the WT background (**Fig. 3A and B**), suggesting that loss of TgGRA17 reduces access to compensatory host-derived metabolites when parasite glycolysis is compromised.

**Figure 3.**
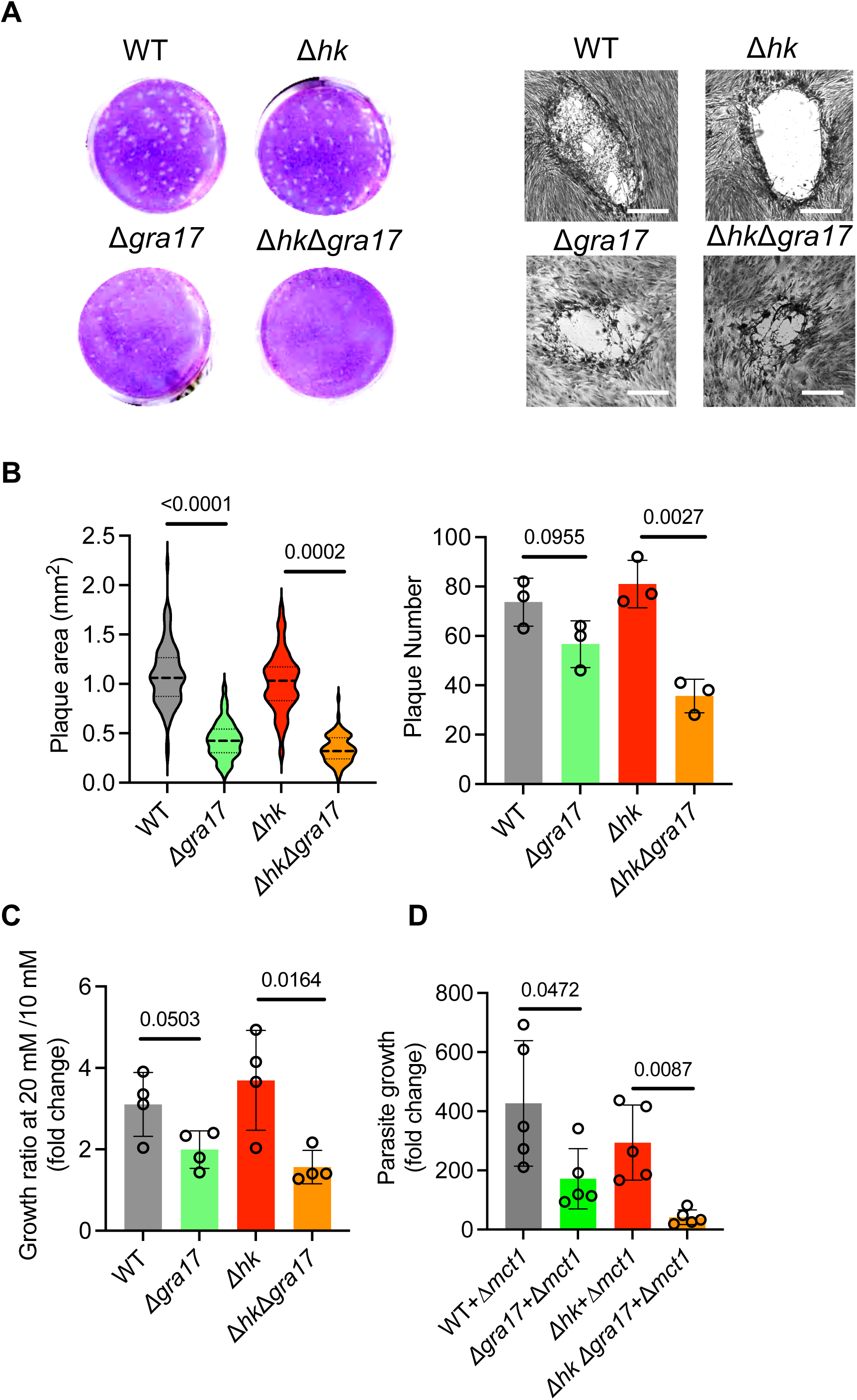
The dense granule protein TgGRA17 promotes uptake of host-derived glycolytic metabolites. (A) Representative plaque assay images of confluent HFF monolayers infected with ∼200 tachyzoites per well and cultured for 7 d prior to crystal violet staining. (B) Plaque area (≥20 plaques per replicate) and plaque number were quantified from 3 independent biological replicates. (C) Enhanced parasite growth at higher lactate concentrations measured by luciferase assay. Parasites were cultured in medium containing 10 mM or 20 mM lactate, and growth is presented as the 20 mM/10 mM ratio for the indicated strains. Data are shown as mean ± SD from 4 independent biological replicates. (D) TgGRA17 disruption in the Δ*hk* background resulted in a significantly greater impairment of host monocarboxylate utilization compared with the WT background. Statistical significance was assessed by two-tailed unpaired Student’s *t*-test (Panel B and C) and two-tailed paired Student’s *t*-test (Panel D).

To more directly test whether TgGRA17 facilitates access to exogenous lactate/pyruvate, we compared growth responses at two substrate concentrations. Parasites (WT, Δ*gra17*, Δ*hk*, Δ*hk*Δ*gra17*) were grown in medium containing 10 mM versus 20 mM lactate, and the ratio of growth (20 mM / 10 mM) was used as a proxy for the extent to which increased lactate availability enhances replication. This ratio decreased by ∼58% in Δ*hk*Δ*gra17* relative to Δ*hk* and by ∼35% in Δ*gra17* relative to WT (**Fig. 3C**). In addition, we cultured Δ*hk* and Δ*hk*Δ*gra17* parasites in HEK293Δ*mct1* cells, which have elevated intracellular monocarboxylates. Similarly, loss of TgGRA17 produced a more pronounced growth defect in the Δ*hk* background than in WT parasites, with growth reduced by 86% versus 60%, respectively (**Fig. 3D**). These data support a model in which TgGRA17-dependent PVM permeability promotes parasite access to host/extracellular monocarboxylates.

### 6. Combined loss of endogenous production and exogenous import of glycolytic metabolites compromises mitochondrial function and ATP production

To determine how endogenous versus exogenous pyruvate acquisition supports mitochondrial physiology, we performed a modified mitochondrial stress assay (24) to measure basal, maximal, and spare capacity OCR (**Fig. 4A**). Basal OCR was similar in Δ*hk* and Δ*fnt1/2/3* parasites compared with WT, whereas Δ*hk*Δ*fnt1/2/3* parasites exhibited a pronounced reduction in basal OCR (**Fig. 4B**). Thus, either endogenous glycolysis or exogenous import is sufficient to sustain basal respiration, but loss of both pathways compromises mitochondrial respiration. Maximal OCR and spare respiratory capacity were severely reduced in Δ*hk* and Δ*hk*Δ*fnt1/2/3* parasites, but not in Δ*fnt1/2/3* parasites (**Fig. 4B**), suggesting that endogenous glycolytic pyruvate is particularly important for meeting increased energetic demand, while imported substrates primarily help maintain baseline function.

**Figure 4.**
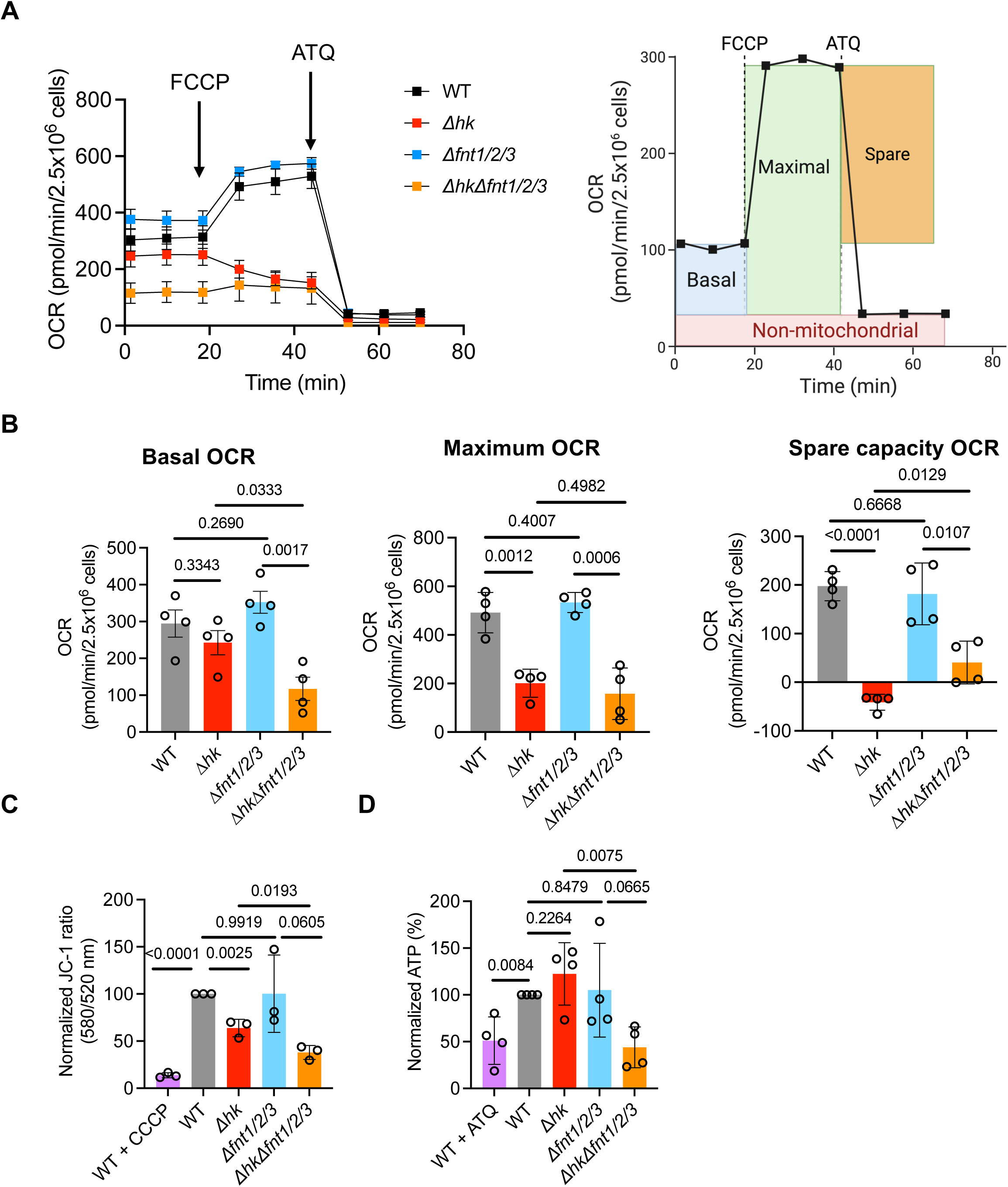
Disruption of pyruvate homeostasis impairs mitochondrial respiration and ATP production. (A) Representative Seahorse OCR assays for the indicated strains. FCCP was injected to elicit maximal respiration, followed by atovaquone (ATQ) to inhibit complex III. (B) Basal respiration, maximal respiration, and spare respiratory capacity quantified from Seahorse mitochondrial stress tests (n = 4 independent biological replicates). (C) Mitochondrial membrane potential measured by JC-1 staining and reported as the red/green fluorescence ratio (emission 585/520 nm; excitation 488 nm). WT parasites treated with 50 μM CCCP served as a positive control for mitochondrial depolarization. (D) Cellular ATP levels quantified by luminescence from parasite lysates and normalized to WT. WT parasites treated with ATQ served as a positive control for reduced ATP production. Data are presented as mean ± SD with 3-4 individual biological replicates indicated in each panel. Statistical significance was assessed by two-tailed unpaired Student’s *t*-test (Panel B, C, and D).

We next assessed mitochondrial membrane potential using JC-1 staining. Compared with WT parasites, Δ*fnt1/2/3* parasites showed no detectable loss of membrane potential. In contrast, Δ*hk* parasites exhibited an ∼30% reduction, and Δ*hk*Δ*fnt1/2/3* parasites displayed a near-complete collapse of membrane potential, comparable to CCCP-treated controls (**Fig. 4C**). These results indicate that endogenously produced pyruvate makes a substantial contribution to mitochondrial respiration and maintenance of the electrochemical gradient, while exogenous monocarboxylate uptake provides a synergistic contribution that becomes critical when endogenous production is impaired. Despite these functional defects, mitochondrial morphology assessed by TgF1β staining appeared normal (**Supplemental Fig. 11**). Total ATP levels paralleled the membrane potential measurements. ATP in Δ*hk*Δ*fnt1/2/3* parasites was reduced by ∼50% relative to WT (**Fig. 4D**). Because pyruvate can also enter the apicoplast, a chloroplast-derived essential organelle in *Toxoplasma*, we stained parasites with TgATrx1 (25), an apicoplast marker, and did not observe noticeable changes in apicoplast morphology (**Supplemental Fig. 12**).

Collectively, these data show that loss of endogenous glycolysis alone has a limited effect on basal respiration, whereas simultaneous disruption of endogenous pyruvate production and exogenous monocarboxylate uptake severely compromises mitochondrial function and cellular energy production. This exogenous monocarboxylate acquisition pathway becomes especially important when parasite glucose metabolism is impaired, for example, under low-glucose conditions *in vivo*, highlighting its role in sustaining mitochondrial activity and overall parasite viability by supplementing lactate/pyruvate metabolism.

## Discussion

Pyruvate is a central metabolic node that links glycolysis to mitochondrial respiration and supports diverse biosynthetic pathways. In most eukaryotic cells, glycolysis is the primary source of pyruvate, and hexokinase (HK) catalyzes the committed first step. Surprisingly, deletion of *TgHK* causes little to no growth defect in nutrient-rich D10% medium (18), indicating that *Toxoplasma gondii* can compensate for reduced glycolytic pyruvate production through alternative metabolic strategies. Consistent with this idea, prior work shows that the parasites under glucose deprivation conditions increase reliance on glutamine entry into the TCA cycle (*e.g.,* via glutaminolysis and/or the GABA shunt) to sustain viability (8, 26). These observations raise a broader question: in addition to endogenous rewiring, does *T. gondii* also exploit exogenous carbon sources to maintain bioenergetic and metabolic homeostasis during infection?

Here, intrigued by this premise, we investigated how *T. gondii* acquires exogenous monocarboxylates, focusing on lactate and pyruvate. By generating a series of TgFNT (formate-nitrite transporter) null mutants, we find that TgFNT1 and TgFNT2 jointly mediate the uptake and utilization of extracellular lactate and pyruvate. In contrast to prior conclusions that loss of TgFNT1 alone is sufficient to block lactate entry for cytoplasm acidification (11), our functional measurements showed that single deletion of TgFNT1 does not abolish lactate import and utilization, indicating substantial redundancy between TgFNT1 and TgFNT2. This interpretation also aligns with findings that TgFNT1 and TgFNT2 can contribute to parasitophorous vacuole (PV) acidification through transport of glycolytic metabolites (13), although such acidification assays reflect export and net proton-coupled flux rather than import *per se*. Notably, TgFNT2 has been reported to be expressed at lower abundance than TgFNT1 (11), yet our data suggest that even relatively low TgFNT2 activity can provide meaningful transport capacity under physiological conditions.

To assess potential compensation between TgFNT1 and TgFNT2, we examined expression in reciprocal deletion backgrounds. TgFNT2 mRNA increased in Δ*hk*Δ*fnt1* by ∼3.5-fold (**Supplemental Fig. 13A**), but TgFNT2 protein levels were not detectably elevated by immunoblot (**Supplemental Fig. 13B**), suggesting post-transcriptional regulation and/or constraints on protein expression and stability. In contrast, we did not observe evidence for transcriptional or translational upregulation of TgFNT1 in Δ*hk*Δ*fnt2* (**Supplemental Fig. 13B**). Because the transport kinetics and substrate affinities of TgFNT1 and TgFNT2 for lactate and pyruvate remain unknown, it remains possible that differences in abundance are offset by differences in catalytic efficiency under relevant substrate concentrations. Direct measurement of kinetic parameters using recombinant transporters reconstituted in proteoliposomes would help resolve their relative contributions and clarify whether TgFNT1 and TgFNT2 operate with similar efficiencies at physiological lactate/pyruvate levels.

A central outcome of this study is that endogenous and exogenous sources of monocarboxylates can be genetically separated to probe their distinct contributions to parasite fitness and virulence. Neither Δ*hk* (endogenous limitation) nor Δ*fnt1/2/3* (exogenous limitation) shows a strong growth defect in rich medium, but combined disruption (Δ*hk*Δ*fnt1/2/3*) produces a drastic reduction of plaque formation and acute virulence. These epistatic relationships support a model in which exogenous monocarboxylate uptake via TgFNTs acts as a supplemental pathway that becomes critical when endogenous glycolytic production is impaired. Importantly, we did not observe evidence that TgFNTs mediate glutamine uptake since OCR responses to glutamine injection in Δ*fnt1/2/3* were indistinguishable from controls (**Supplemental Fig. 14**), arguing against an indirect effect of TgFNT loss on the major compensatory glutamine route. Together, these results support the conclusion that TgFNT-dependent import of lactate/pyruvate helps maintain mitochondrial function by supplying oxidizable substrates, particularly when glycolytic pyruvate production is reduced.

Our data further indicate that lactate and pyruvate are not functionally equivalent inputs. Because lactate must be converted to pyruvate for downstream metabolism, Δ*ldh1* parasites exhibit impaired lactate-supported bioenergetics compared with pyruvate. *In vivo*, this distinction is likely important because circulating glucose concentrations in healthy individuals (∼3.9–5.6 mM) are far lower than the glucose level in D10 medium (25 mM) (27). In addition, lactate is typically present at substantially higher concentrations than pyruvate in mammalian tissues (approximately 0.5–20 mM versus ∼100 µM, respectively) (28, 29), and lactate levels can vary widely across physiological and pathological contexts, including inflammation, hypoxia, and conditions with high glycolytic flux. The attenuated virulence observed in Δ*fnt1/2* and Δ*fnt1/2/3* mutants relative to WT supports the idea that host-derived monocarboxylates contribute to acute infection, with the strongest defects emerging when uptake of both lactate and pyruvate is restricted. The comparatively modest attenuation of Δ*ldh1* may reflect continued access to exogenous pyruvate even when lactate conversion is blocked, consistent with a model in which both monocarboxylates can contribute, but lactate may represent a major abundant input *in vivo*. Previous work reported that a type II ME49Δ*ldh1* strain failed to cause lethal infection in mice when animals were inoculated with 100 tachyzoites (30), a dose that is typically sublethal for type II parasites, suggesting that incorporation of exogenous monocarboxylates may be particularly important in type II strains. Future animal studies spanning a range of inocula and strain backgrounds (*e.g.,* RH, ME49, or VEG) will enable a quantitative comparison of lactate-specific defects with more general monocarboxylate uptake impairments.

These findings also fit into a broader evolutionary context. FNT family transporters are widespread across prokaryotes and eukaryotes (31). *T. gondii* encodes three FNTs (TgFNT1-3) that can mediate bidirectional transport of monocarboxylates (11–13), whereas *Plasmodium falciparum* encodes a single essential PfFNT, characterized primarily for exporting excess lactate produced during the highly glycolytic trophozoite stage (32). The differing essentiality of FNTs in *P. falciparum* versus *T. gondii* likely reflects major differences in carbon metabolism and the extent of glycolytic reliance across species and life stages. In *P. falciparum*, PfFNT is essential during the trophozoite stage, consistent with the parasite’s heavy dependence on glycolysis and the need to export lactate to avoid toxic accumulation (33, 34). In contrast, FNTs are dispensable in *T. gondii* tachyzoites under standard nutrient-rich culture conditions (11). Notably, PfFNT becomes dispensable in *Plasmodium* gametocytes, which exhibit greater TCA-cycle activity that may reduce lactate production and lessen the requirement for lactate export (34). In *Toxoplasma* tachyzoites, FNTs are dispensable in nutrient-rich conditions, but our results indicate that they provide a physiologically important advantage *in vivo*, where nutrient availability and host metabolite composition differ strikingly from culture media.

An additional layer that must be traversed for host nutrient access is the parasitophorous vacuole membrane (PVM), which must permit small metabolites to reach the parasite. Prior work identifies several dense granule proteins (*e.g.,* TgGRA17 and TgGRA23) that contribute to PVM small-solute permeability (4). Because lactate and pyruvate fall within the size range expected to traverse PVM pores, we hypothesized that restricting PVM permeability would further sensitize parasites with compromised glycolysis. Consistent with this model, *TgGRA17* deletion in the Δ*hk* background decreased plaque size and plaque number by 67% and 56%, respectively, compared with decreases of 60% and 23% in the WT background (**Fig. 3A–B**), supporting the idea that loss of TgGRA17 reduces access to compensatory host-derived metabolites when glycolytic production is limited. Although the substrate specificity of PVM pores remains incompletely defined, the genetic interaction between Δ*hk* and Δ*gra17* is consistent with a TgGRA17-mediated import route across the PVM that delivers host-derived metabolites to the parasite and supports fitness under metabolic constraint. Because loss of both TgGRA17 and TgGRA23 is lethal (4), inducible knockdown of TgGRA17 in a TgGRA23-deficient background would provide a powerful approach to test whether near-complete restriction of PVM permeability abolishes access to host-derived monocarboxylates during intracellular growth.

Our data support a two-barrier, two-input model for monocarboxylate supply during intracellular growth (**Fig. 5**). Host-derived lactate and pyruvate in the cytosol first traverse the parasitophorous vacuole membrane (PVM) through TgGRA17-dependent small-solute permeability and then cross the parasite plasma membrane via TgFNT1/2. Once inside the parasite, lactate requires LDH1 to be converted to pyruvate, providing oxidizable substrate for mitochondrial respiration. When endogenous glycolytic pyruvate production is intact, loss of FNT-mediated uptake has limited impact in nutrient-rich conditions; however, when endogenous production is constrained (Δ*hk*), the parasite becomes increasingly dependent on this import route to usurp host monocarboxylates, and dual disruption (Δ*hk*Δ*fnt1/2/3*) results in profound mitochondrial depolarization and reduced ATP.

**Figure 5.**
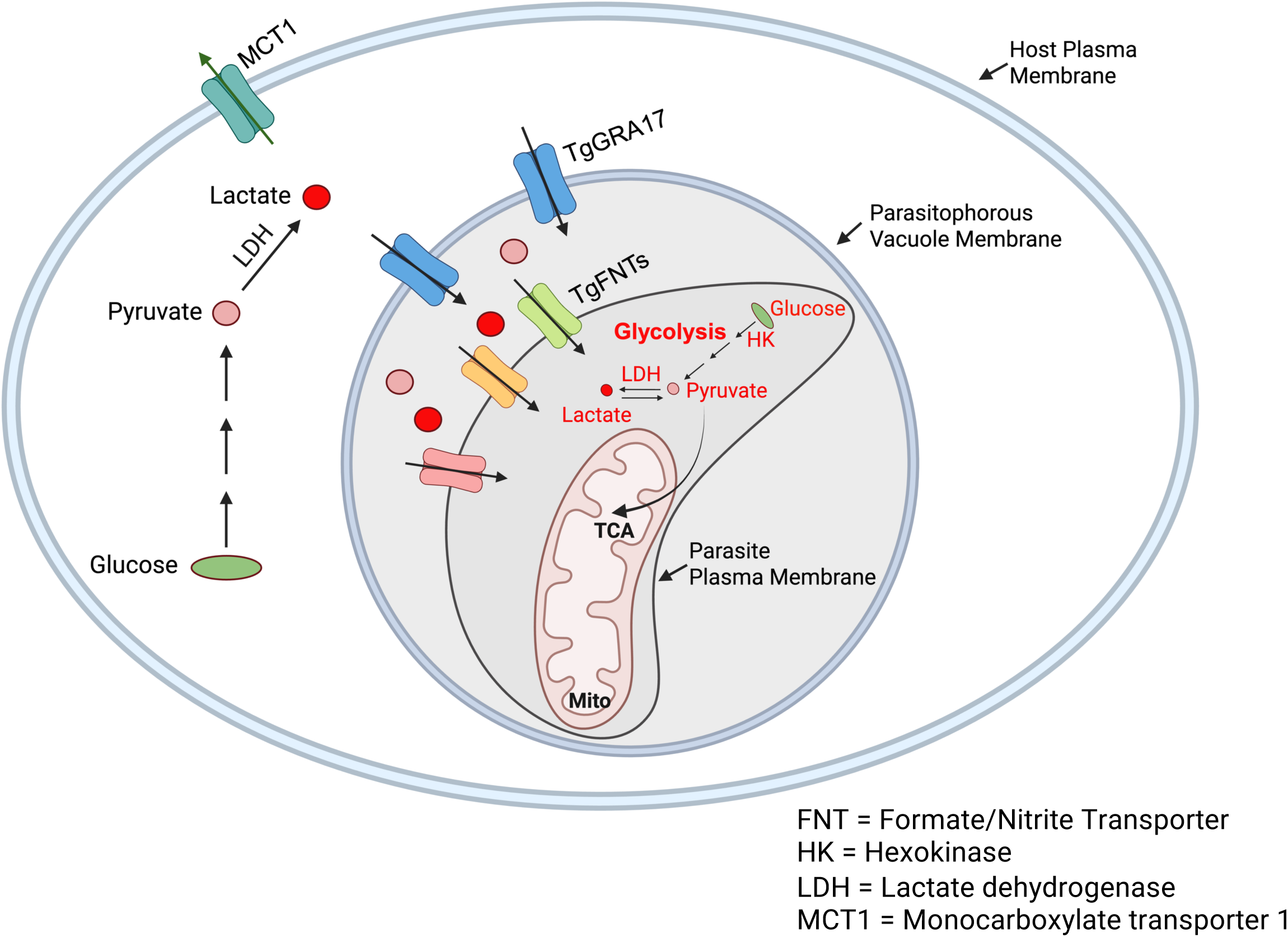
A two-step model for incorporation of exogenous monocarboxylates by *Toxoplasma gondii.* See the main text for a detailed description.

Together, these findings indicate that *T. gondii* maintains mitochondrial bioenergetic capacity by drawing on two complementary carbon sources: internally generated glycolytic pyruvate and host-derived monocarboxylates imported across the PVM and parasite plasma membrane. This two-step acquisition route depends on TgGRA17-mediated permeability of the PVM and subsequent TgFNT-mediated transport across the parasite plasma membrane, providing metabolic flexibility during infection when nutrient availability differs from culture conditions. Because simultaneous impairment of endogenous production and exogenous uptake triggers a sharp bioenergetic collapse, this pathway represents a potential vulnerability that could be exploited for therapeutic intervention against toxoplasmosis.

## Materials and Methods

### Ethics statement

All animal procedures were conducted in compliance with the Public Health Service Policy on Humane Care and Use of Laboratory Animals and AAALAC guidelines and were approved by the Clemson University Institutional Animal Care and Use Committee (Animal Welfare Assurance A3737-01; protocol AUP2016-012). Mice meeting moribund criteria were euthanized by CO_2_ inhalation in accordance with AVMA Guidelines for the Euthanasia of Animals.

### Chemicals and reagents

Oligonucleotide primers were purchased from Eurofins (**Supplemental Table 1**). The chemicals, essential antibodies, plasmids, equipment, and software are listed in **Supplemental Table 2.** Unless otherwise noted, routinely used chemicals were analytical grade and purchased from VWR.

### Host cell and parasite culture

Human foreskin fibroblasts (HFFs; ATCC SCRC-1041) were maintained at 37°C in 5% CO_2_ in DMEM (4.5 g/L glucose; GenDepot) supplemented with 10% fetal bovine serum (FBS; Cytiva), 10 mM HEPES, and an additional 2 mM L-glutamine (4 mM in total). The parental *Toxoplasma* gondii RHΔ*ku80*Δ*hxg* strain was provided by the Carruthers laboratory and maintained by serial passage in confluent HFF monolayers every 48 h (35). The Δ*gra17* strain was provided by the Saeij laboratory and has been described previously (4).

### Generation of transgenic *Toxoplasma* strains

#### Transfection by electroporation

Freshly egressed tachyzoites were released from host cells by syringe passage and separated from host debris by filtration through a 3-µm polycarbonate filter. Parasites were washed and resuspended in Cytomix buffer (10 mM KPO_4_, pH 7.6; 120 mM KCl; 5 mM MgCl_2_; 0.15 mM CaCl_2_; 25 mM HEPES; 2 mM EGTA, pH 7.6). Parasites were pelleted at 1,000 *× g*, resuspended in 400 µL Cytomix, and transferred to a 4-mm electroporation cuvette. Cas9/sgRNA-expressing plasmid DNA and repair template DNA were combined in Cytomix (typically a 5:1 repair-template-to-plasmid mass ratio) to a final volume of 100 µL and mixed with the parasite suspension in a total volume of 500 µL. Parasites were electroporated using two 1.5-kV pulses and immediately transferred onto confluent HFF monolayers in complete D10% medium.

#### Gene disruption by CRISPR/Cas9

Targeted knockouts (TgHK, TgFNT1, TgFNT2, and TgFNT3) were generated using CRISPR/Cas9-mediated double-strand breaks with homology-directed repair as described before (36, 37). Repair templates were amplified to include ∼50-bp homology arms flanking the targeted locus and a selectable marker cassette. Transfected populations were selected with the appropriate drug, cloned by limiting dilution, and verified by diagnostic PCR using primers spanning the coding region and both integration junctions (**Supplemental Table 1**).

#### Epitope tagging

For C-terminal tagging, RHΔ*ku80*Δ*hxg* and RHΔk*u80*Δ*hxg*Δ*hk* parasites were transfected with a Cas9/sgRNA plasmid targeting the 3′ end of the gene of interest and a repair template encoding the epitope tag with ∼50-bp flanking homology arms. Drug-resistant populations were cloned by limiting dilution, and correct integration was confirmed by junction PCR (**Supplemental Table 2**).

#### NanoLuc reporter strains

NanoLuciferase (NLuc) reporter parasites were generated by electroporation of an NLuc expression plasmid, followed by drug selection and cloning by limiting dilution. Clones were screened for luminescence output, and those with robust signal were used for growth assays.

### Generation of transgenic host cell strains

#### MCT1 knockout in HEK293 cells

HEK293 cells were transfected using JetOptimus (Polyplus) with 3 Cas9/sgRNA plasmids targeting the first exon of human SLC16A1 (MCT1) in combination. Edited populations were cloned after 3 rounds of passage. Clones were validated by genotyping.

#### Transient expression of the Laconic lactate sensor

The Laconic FRET lactate sensor plasmid was obtained from Addgene (Plasmid #: 44238) and has been described previously (22). HEK293 and HEK293Δ*mct1* cells were seeded in black, clear-bottom 96-well plates and transfected with Laconic plasmid DNA using JetOptimus according to the manufacturer’s protocol. Cells were analyzed 24 h after transfection.

#### RT-qPCR

Parasites were harvested from infected HFF monolayers, separated from host debris by 3-µm filtration, and washed in cold 1x PBS. Total RNA was extracted using the Quick-RNA Miniprep Kit (Zymo Research) according to the manufacturer’s instructions. Relative transcript abundance was quantified using the Luna Universal One-Step RT-qPCR kit (NEB) with ∼200 ng RNA per reaction. TgActin was used as an internal reference gene. Relative expression was calculated by the ΔΔC_T_ method and reported relative to the parental strain.

#### SDS-PAGE and immunoblotting

Freshly egressed parasites were purified by 3-µm filtration, washed in cold 1x PBS, and counted. Parasites were resuspended in SDS sample buffer (40 mM Tris-HCl, pH 6.8; 1% (m/v) SDS; 5% (v/v) glycerol; 50 mM DTT; 0.0003% (m/v) bromophenol blue) at 5 × 10^8^ tachyzoites/mL and incubated at 98°C for 5 min or 37 °C for 20 min. Low temperature incubation was applied to transmembrane proteins to avoid precipitation. Proteins were separated by SDS-PAGE and transferred to PVDF membranes using semi-dry transfer. Membranes were blocked in 5% nonfat milk in PBS-T (0.1% Tween-20), incubated with primary antibodies in 1% milk/PBS-T, and then with HRP-conjugated secondary antibodies. Signal was developed using SuperSignal West Pico (Thermo Fisher Scientific) and imaged on an Azure C600 system.

#### Immunofluorescence microscopy

Purified tachyzoites were added to confluent HFF monolayers on chamber slides and incubated for 1 h (pulse-invaded) or 24 h (replicating). Samples were fixed and processed by standard immunofluorescence staining. Images were acquired on a Leica DMi8 inverted epifluorescence microscope and processed using Leica LAS X.

#### Plaque assay

Freshly egressed parasites were purified by 3-µm filtration and pelleted at 1,000 *× g* for 10 min. Parasites were resuspended in complete medium at 100 tachyzoites/mL, and 200 tachyzoites were added to each well of a confluent six-well plate of HFFs. Plates were incubated for 7 d, fixed, and stained with 0.02% crystal violet. Plaques were imaged with a Leica DMi8 microscope at 25*×* magnification, and individual plaques areas were quantified in Fiji and plotted for comparison.

### Luciferase-based growth assays

#### Growth in HEK293 cells

HEK293 cells were seeded in white 96-well plates and grown in D10% medium for 48 h prior to infection. NLuc-expressing parasites were added at 1 × 10^4^ tachyzoites/mL (150 µL per well). After 4 h, wells were washed and switched to low-serum medium (D1% medium) to limit host cell overgrowth. Luminescence was measured every 24 h for 96 h and normalized to the 4-h time point as published before (38).

#### Growth in HFFs under test media

Confluent HFF monolayers in white 96-well plates were infected with NLuc parasites for 4 h, washed, and switched to phenol red-free base medium containing 10% dialyzed serum and the indicated nutrients as specified in the figure legends. Luminescence was measured every 24 h for 48 h.

#### Murine infection model

Outbred CD-1 mice were infected subcutaneously with 100 tachyzoites in 1x PBS and monitored daily for 30 d. Moribund mice were euthanized in accordance with IACUC-approved humane endpoint criteria. Surviving mice were assessed for seroconversion by ELISA using *Toxoplasma* antigen-coated plates. Survivors were rested for 10 d and challenged with 1,000 parental-strain tachyzoites. Survival curves were compared using the log-rank (Mantel-Cox) test.

### Seahorse oxygen consumption rate assays

#### Mito stress test

A 24-well Seahorse XF plate was coated with Cell-Tak (Corning) per the manufacturer’s instructions, and the sensor cartridge was hydrated according to the manufacturer’s protocol. Purified parasites were resuspended in Seahorse XF DMEM assay medium supplemented with 10 mM glucose and 4 mM glutamine at 2.5 × 10^7^ tachyzoites/mL, and 100 µL was seeded per well. Plates were centrifuged at low speed to promote even attachment, which was confirmed by light microscopy. FCCP (2 µM) and atovaquone (1 µM) were injected as indicated, following established *Toxoplasma* Seahorse workflows (24).

#### Substrate-stimulated OCR

Baseline OCR was recorded in assay medium (10 mM glucose, 4 mM glutamine) before acute injection with assay medium alone or with lactate or pyruvate (10 mM final). Percent change was calculated relative to baseline. Data were analyzed in GraphPad Prism using an unpaired two-tailed Student’s *t*-test.

#### Laconic FRET measurements

At ∼24 h post-transfection, wild type HEK293 and HEK293Δ*mct1*cells plated in 96-well black opaque plates were washed and measured in 1x PBS using a BioTek H1 Hybrid plate reader at excitation 430 nm and emissions 480 and 535 nm for mTFP and Venus, respectively (5). The mTFP/Venus ratio was background-corrected using control-transfected wells and analyzed in GraphPad Prism and statistical significance was calculated using student’s paired *t*-test.

#### Lactate secretion assay

Parasites were resuspended at 5 × 10^7^ tachyzoites/mL in phenol red-free DMEM containing 10 mM glucose and incubated for 1 h. Lactate in supernatants was quantified using a coupled enzymatic assay containing 0.5 U/mL diaphorase, 3.3 mM NAD⁺, 3.3 U/mL LDH, and 11.1 µM resazurin in 50 mM KPO₄ buffer. Supernatant (20 µL) was mixed with reaction buffer (180 µL) and incubated for 30 min at room temperature in 96-well clear, flat-bottomed plates. Fluorescence was measured (Ex 535 nm/Em 590 nm). Lactate concentrations were calculated using a sodium L-lactate standard curve.

#### ATP quantification

Purified parasites were washed in cold 1x PBS and resuspended at 2 × 10^7^ tachyzoites/mL. Parasite suspension (50 µL) was dispensed into white 96-well plates, and ATPlite reagent (Revvity; 100 µL) was added. Luminescence was recorded, averaged across at least 3 technical replicates, and normalized to parental-strain controls.

#### JC-1 mitochondrial membrane potential assay

Purified parasites were resuspended at 1 × 10^8^ tachyzoites/mL in warm PBS and stained with 2 µM JC-1 for 15 min at 37 °C. As a depolarized control, parasites were pre-incubated with CCCP (50 µM) for 5 min prior to JC-1 staining. Parasites were washed once in warm 1x PBS and analyzed in black 96-well plates (green form: Ex 488/Em 520; red form: Ex 488/Em 585). The red/green ratio was background-corrected using unstained controls.

### *Toxoplasma* Replication assay

Purified parasites were inoculated onto confluent HFF monolayers in chamber slides and incubated for 24 h before fixation. Samples were stained with anti-TgGRA7 (1:2,000) and DAPI to recognize individual parasitophorous vacuoles (PVs) and parasite nuclei, respectively. Parasites per PV were counted for 100 vacuoles per strain.

### *Toxoplasma* Invasion Assay

Freshly egressed parasites were filter-purified and resuspended in invasion medium (DMEM supplemented with 3% Cosmic Calf Serum). Parasites (4.5 × 10^6^ per well) were added to clear 96-well plates pre-seeded with HFFs in three technical replicates per strain. After 30 min at 37 °C, wells were fixed with 4% formaldehyde for 20 min. To distinguish attached from invaded parasites, wells were first stained under non-permeabilizing conditions with mouse anti-TgSAG1 (1:2000, 1 h), followed by goat anti-mouse IgG-Alexa Fluor 594 (1:1000; Invitrogen) to label extracellular/attached parasites. Wells were then permeabilized with 0.1% Triton X-100 for 10 min and stained with rabbit anti-TgMIC5 (1:1000) followed by goat anti-rabbit IgG-Alexa Fluor 488 (1:1000; Invitrogen) to label total parasites (attached + invaded). DAPI was included to stain host nuclei. For each well, 25 fields of view were acquired at 200 x total magnification using a BioTek Cytation C10 Confocal Imaging Reader and quantified with Cell Count analysis in Agilent BioTek Gen5 software. Invasion efficiency was calculated by using the following equation: (total parasites [green] − extracellular parasites [red]) / total host nuclei (DAPI). Each assay was performed in three independent biological replicates.

### *Toxoplasma* Egress Assay

Parasite egress was quantified using host-cell lactate dehydrogenase (LDH) release. Freshly egressed parasites were filter-purified, resuspended in D10 medium at 5 × 10^5^ parasites/mL, and inoculated into 96-well plates pre-seeded with HFFs at 100 µL per well with three technical replicates per strain. After incubation at 37 °C and 5% CO₂ for 18-24 h, wells were washed and incubated with 50 µL Ringer’s buffer (10 mM HEPES, pH 7.2; 3 mM NaH₂PO₄; 1 mM MgCl₂; 2 mM CaCl₂; 3 mM KCl; 115 mM NaCl; 10 mM glucose; 1% FBS) for 20 min. Egress was induced by adding an equal volume of 1 mM Zaprinast prepared in Ringer’s buffer and incubating for 5 min at 37 °C and 5% CO₂. Uninfected wells treated with Ringer’s buffer containing 1% Triton X-100 or Ringer’s buffer alone served as positive and negative controls, respectively. Supernatants were clarified by centrifugation at 1,000 x g for 5 min twice to remove insoluble debris. LDH activity was measured from 50 µL clarified supernatant using a standard LDH release assay as described previously (37). Egress efficiency was calculated as the following equation: (LDH [sample] – LDH [negative]) / (LDH [positive] − LDH_[negative]).

## Supporting information

Supplemental Table 1 primers

Supplemental Table 2 Resources

## Acknowledgments

We thank our colleagues Drs. Vern Carruthers, Peter Bradley, and David Sibley for generously supplying essential reagents for this study. Funding for this research was provided by the National Institutes of Health grant R01AI143707 (awarded to Z.D.).

## Supplemental Figure and Table Legends

**Supplemental Figure 1. WT and Δ*hk* parasites can utilize exogenous pyruvate to stimulate mitochondrial respiration.** (A) Representative Seahorse OCR traces following acute injection of base medium alone or base medium supplemented with 10 mM pyruvate. (B) Pyruvate-stimulated OCR response quantified as percent increase over the pre-injection basal OCR. Data are shown as mean ± SD from three independent biological replicates. Statistical significance was assessed by two-tailed unpaired Student’s *t*-test.

**Supplemental Figure 2. Generation of parasites lacking hexokinase (HK) and lactate dehydrogenase (LDH1).** (A-C) Diagnostic PCRs confirming replacement of TgHK and/or TgLDH1 with a drug-resistance cassette and correct integration at the 5′ and 3′ junctions (5′ and 3′ arms).

**Supplemental Figure 3. Δ*ldh1* parasites exhibit reduced lactate secretion.** Purified extracellular WT::*NLuc* and Δ*ldh1*::*NLuc* tachyzoites were incubated for 1 h in DMEM containing 10 mM glucose, after which lactate released into the medium was quantified. Statistical significance was assessed using a two-tailed unpaired Student’s *t*-test.

**Supplemental Figure 4. HsMCT1-deficient HEK293 cells exhibit reduced lactate export and elevated cytosolic lactate.** (A) Immunoblot confirmation of HsMCT1 loss in Δ*mct1* HEK293 cells generated in this study. (B) Lactate secretion is reduced in Δ*mct1* HEK293 cells. (C) Cytosolic lactate measured using the Laconic FRET sensor. Representative images show expression of mTFP and Venus fluorescent proteins in cytosol, and the corresponding mTFP/Venus ratio was normalized to WT HEK293 cells and plotted. Data are shown as mean ± SD from 3 independent biological replicates. Statistical significance was assessed by two-tailed unpaired Student’s *t*-test.

**Supplemental Figure 5. Generation of combined TgFNT-deficient parasites in the Δ*hk* background.** (A-C) Diagnostic PCRs confirming loss of the indicated TgFNT coding sequences.

**Supplemental Figure 6. Generation of single TgFNT knockouts in the Δ*hk* background.** (A-B) Diagnostic PCRs confirming replacement of the indicated TgFNT coding sequences with a drug-resistance cassette and correct integration at the 5′ and 3′ junctions (5′ and 3′ arms).

**Supplemental Figure 7. Single ablation of FNT transporters does not abolish lactate- or pyruvate-stimulated respiration.** (A, B) Percent increase in OCR following acute injection of base medium alone or base medium supplemented with 10 mM lactate or 10 mM pyruvate, measured by Seahorse assay for the indicated single-FNT mutant strains (A) and combined-FNT mutant strains (B). Data are shown as mean ± SD from 3 independent biological replicates. Statistical significance was assessed by two-tailed paired Student’s *t*-test.

**Supplemental Figure 8. Generation of single and combined TgFNT knockouts in the WT background.** (A-C) Diagnostic PCRs confirming replacement of the indicated TgFNT coding sequences with a drug-resistance cassette and correct integration at the 5′ and 3′ junctions (5′ and 3′ arms).

**Supplemental Figure 9. Additional phenotypic characterization of TgFNT-deficient strains.** (A) Plaque area and (B) plaque number quantification. (C) Host-cell invasion assay. (D) Intracellular replication assay. (E) Egress assay. Statistical significance was assessed by two-tailed unpaired Student’s *t*-test.

**Supplemental Figure 10. Genetic ablation of TgHK in the Δ*gra17* background.** Diagnostic PCR confirming loss of the TgHK coding sequence in parasites derived from the Δ*gra17* parental strain (provided by Jeroen Saeij, University of California, Davis).

**Supplemental Figure 11. Mitochondrial morphology is preserved despite perturbation of pyruvate homeostasis.** Intracellular parasites were cultured in HFF monolayers for 24 h, fixed, and stained with anti-F1β to visualize the parasite mitochondrion. All strains exhibited the characteristic lasso-shaped mitochondrial morphology. Scale bar, 2 μm.

**Supplemental Figure 12. Apicoplast morphology is unchanged in parasites lacking hexokinase or FNT transporters.** Intracellular parasites were cultured in HFF monolayers for 24 h, fixed, and stained with anti-Atrx1 to label the apicoplast. Parasite nuclei were counterstained with DAPI. No defects in apicoplast biogenesis were observed in the indicated strains. Scale bar, 2 μm.

**Supplemental Figure 13. TgFNT2 transcripts increase in the absence of TgFNT1 without a corresponding rise in protein abundance.** (A) RT-qPCR analysis of TgFNT1 and TgFNT2 transcript levels, shown as fold change relative to the corresponding parental strain. (B) Representative immunoblots of 3×HA-tagged FNT1 and 3×myc-tagged FNT2 from total parasite lysates with actin as a loading control. Densitometry quantification of FNT1-3×HA and FNT2-3×myc levels is normalized to the tagged WT controls. Data are shown as mean ± SD from 3 independent biological replicates. Statistical significance was assessed by two-tailed unpaired Student’s *t*-test.

**Supplemental Figure 14. Glutamine-stimulated respiration is not altered in FNT-deficient parasites.** Percent increase in OCR following acute injection of base medium (DMEM + 10 mM glucose) or base medium supplemented with 10 mM L-glutamine, measured by Seahorse assay for the indicated strains. Data are shown as mean ± SEM (n = 4 independent biological replicates). Statistical significance was assessed by two-tailed paired Student’s *t*-test.

**Supplemental Table 1. Chemicals, antibodies, plasmids, equipment, and software used in this study.**

**Supplemental Table 2. Primers used in this study.**

## Notes

### Competing Interest Statement

The authors have declared no competing interest.

