## Supplemental figures for "A two-membrane gateway for monocarboxylates couples host glycolysis to *Toxoplasma* mitochondrial fitness and virulence"

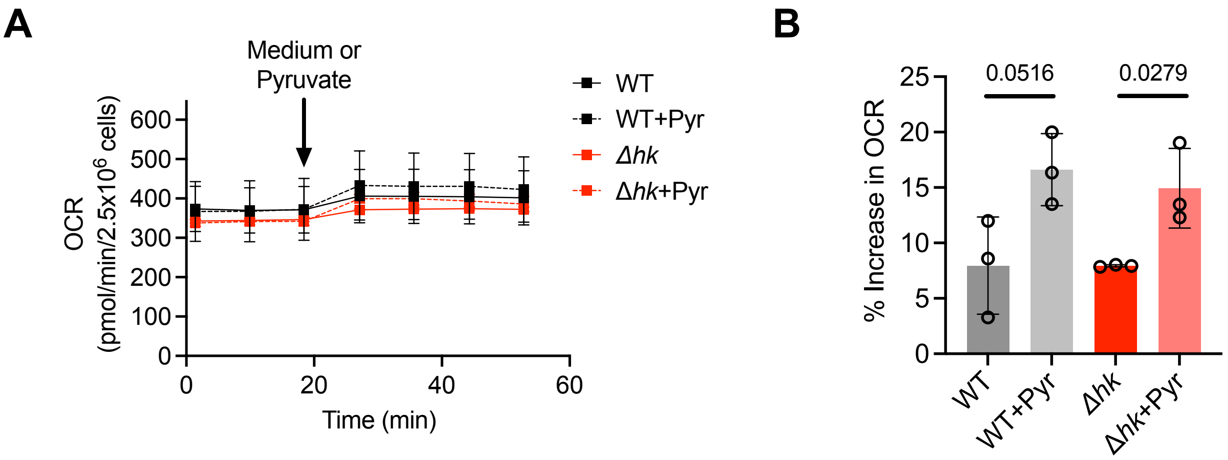


**Supplemental Figure 1**

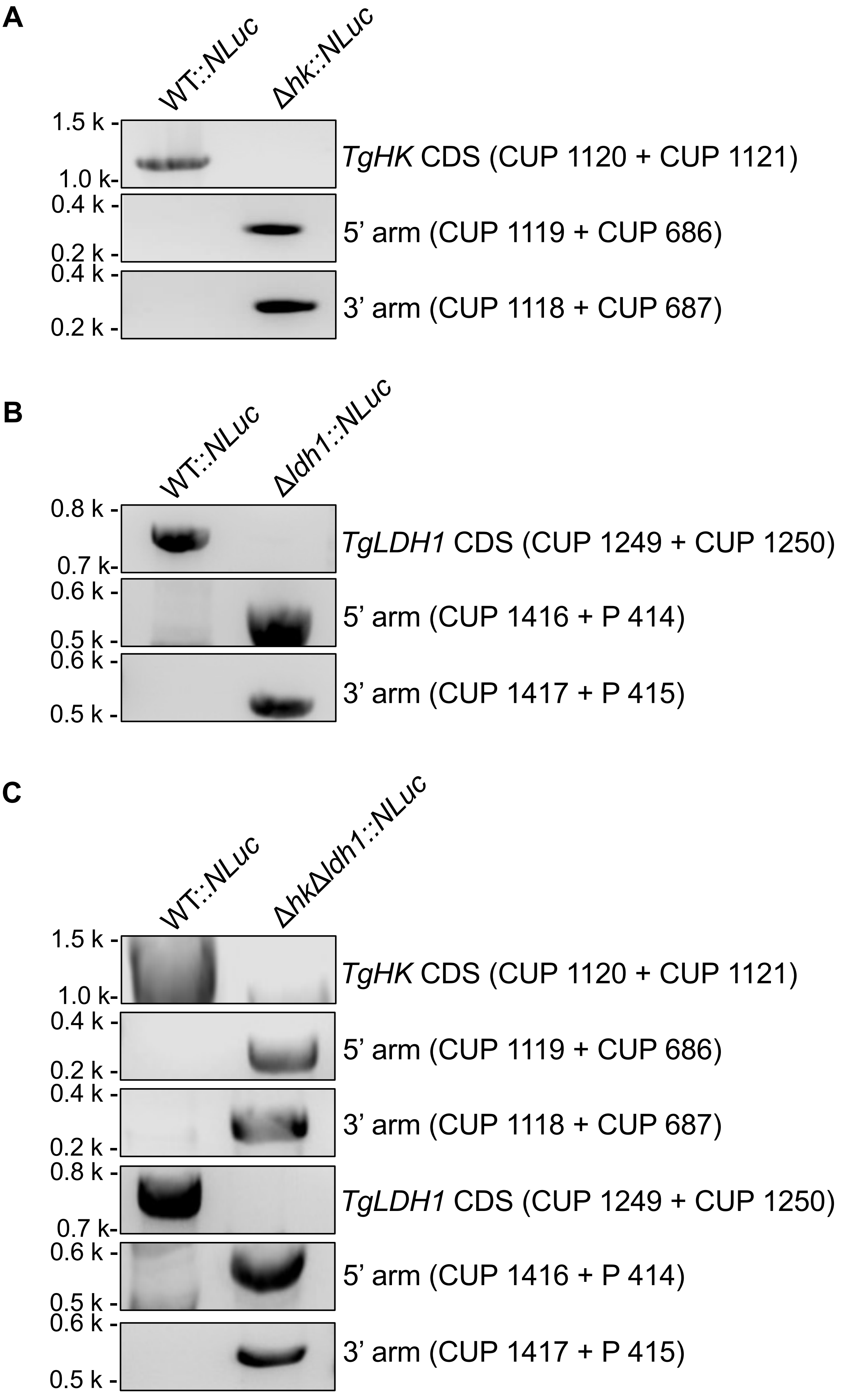


**Supplemental Figure 2**

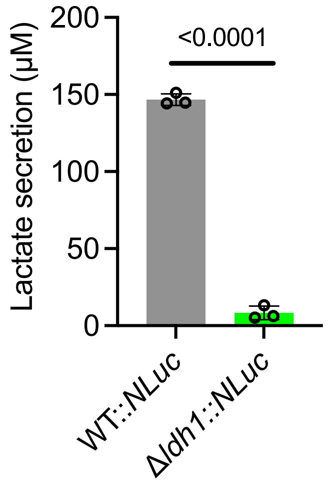


**Supplemental Figure 3**

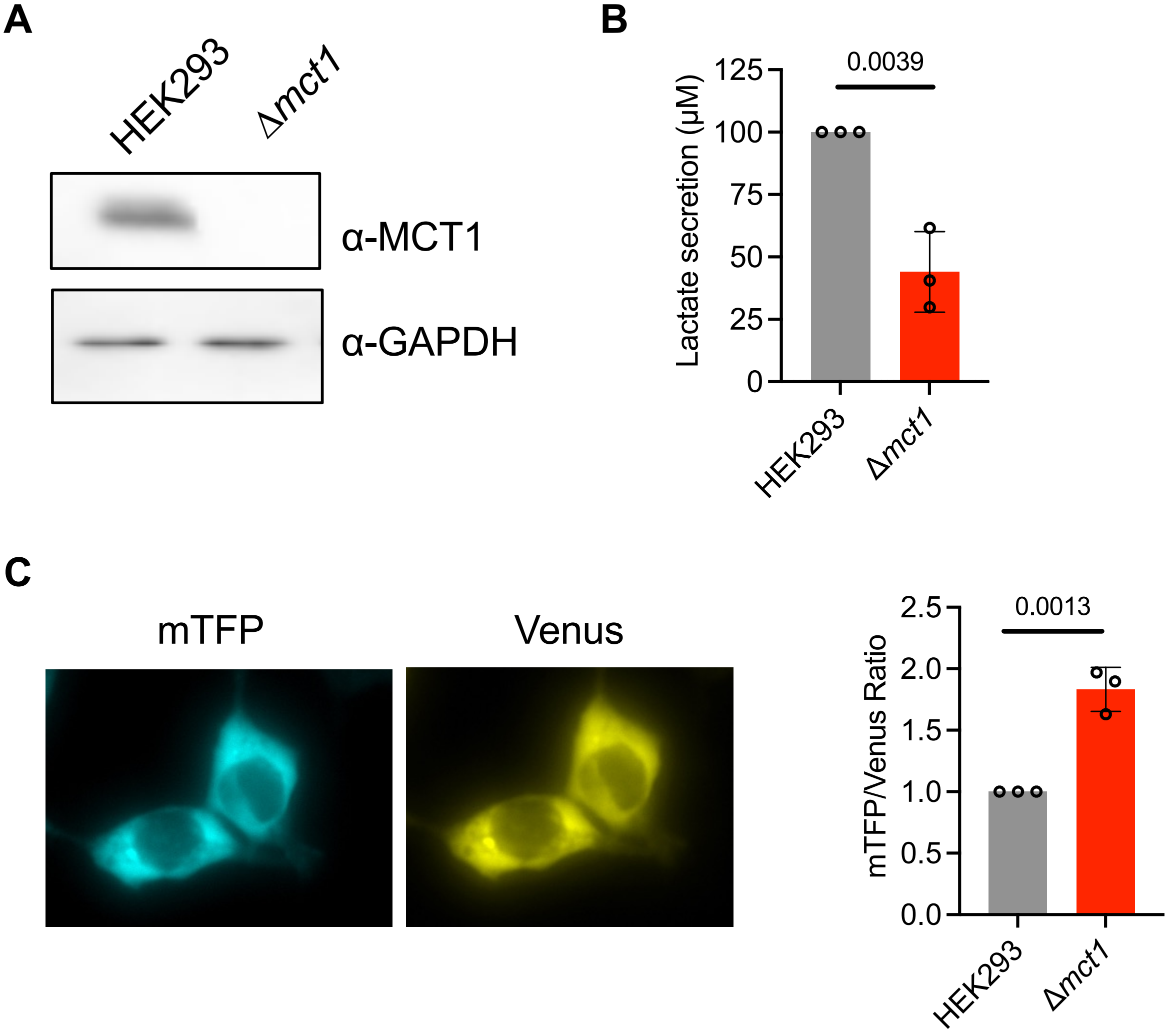


**Supplemental Figure 4**

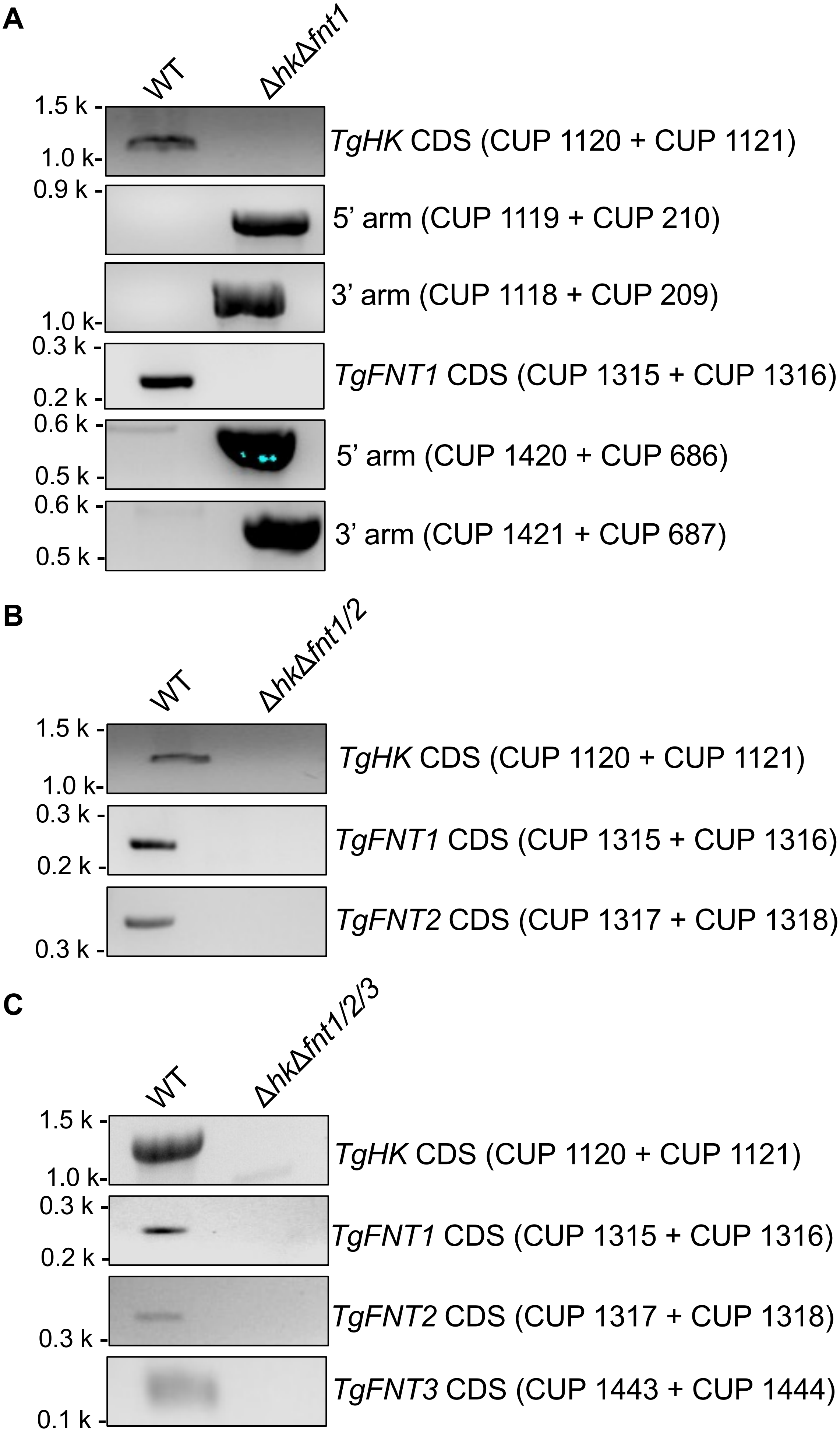


**Supplemental Figure 5**

**A**

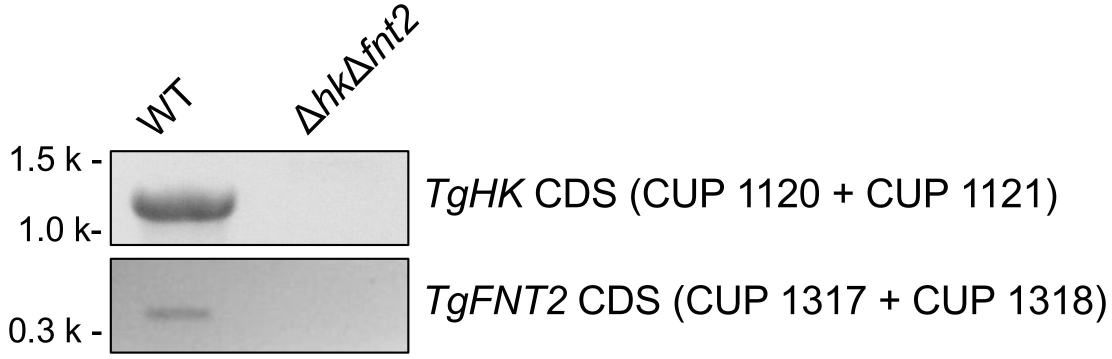


**B**

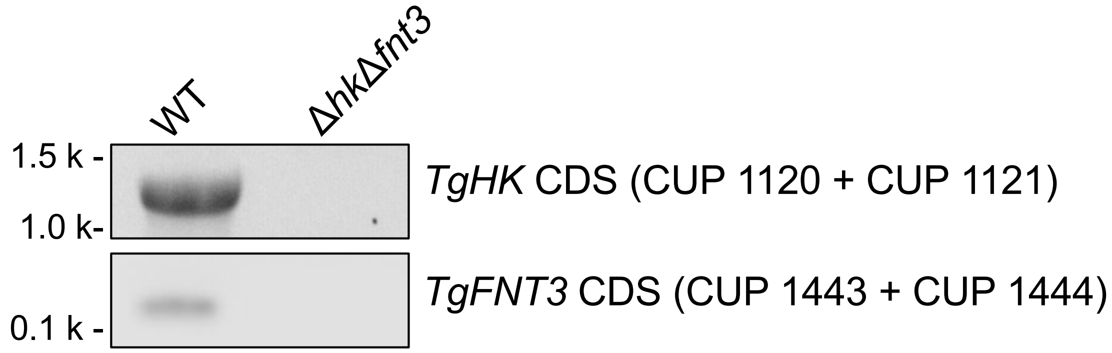


**Supplemental Figure 6**

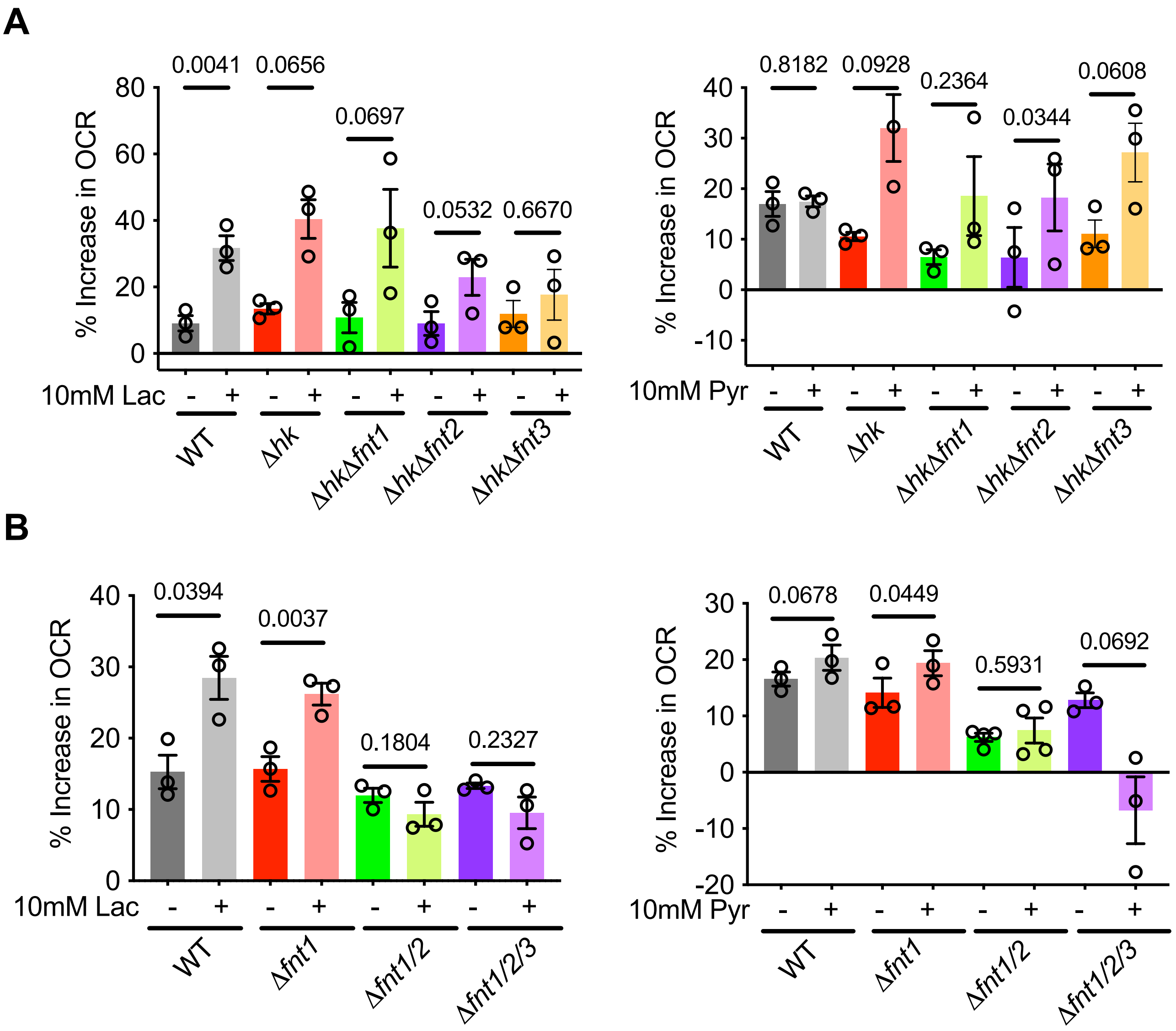


**Supplemental Figure 7**

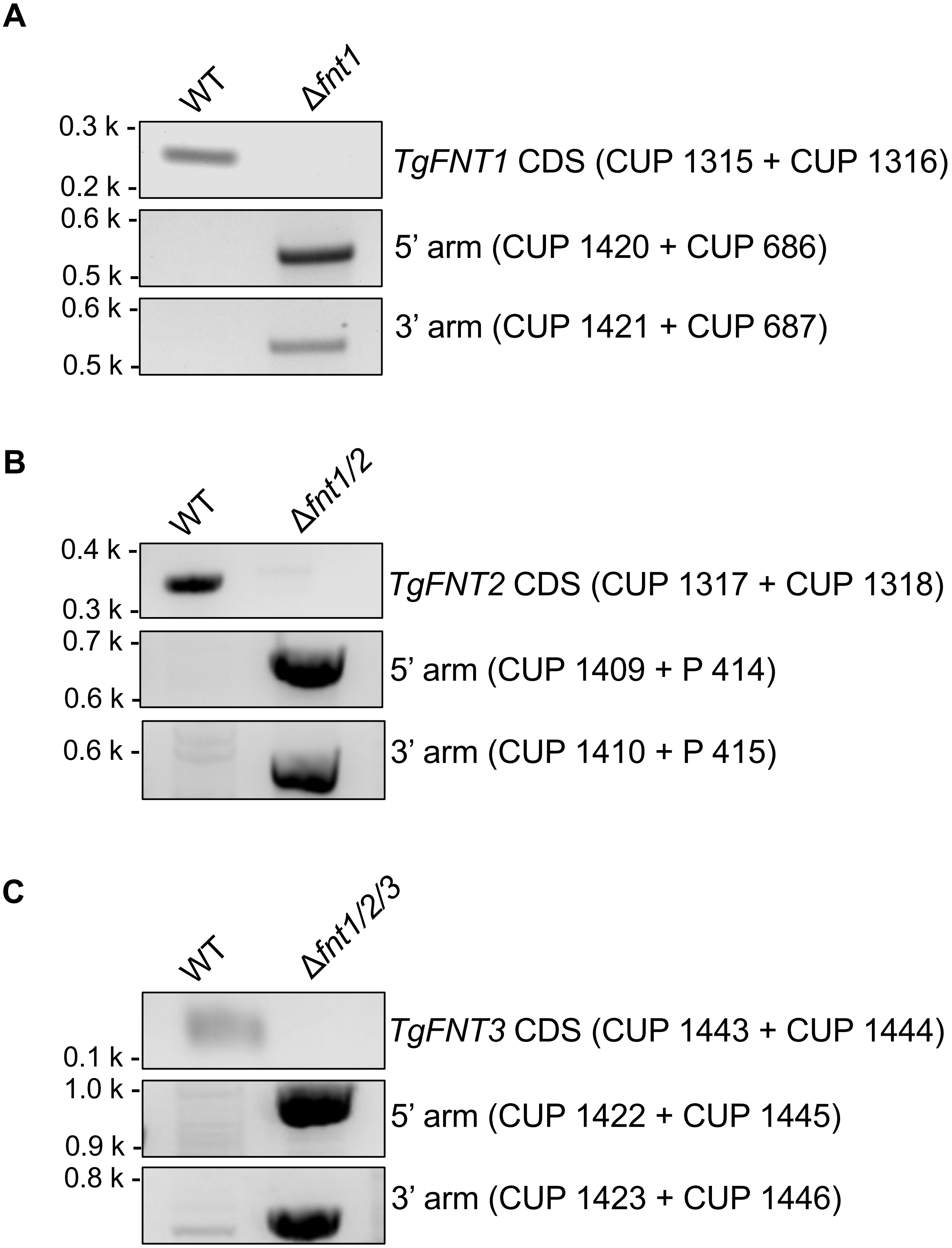


**Supplemental Figure 8**

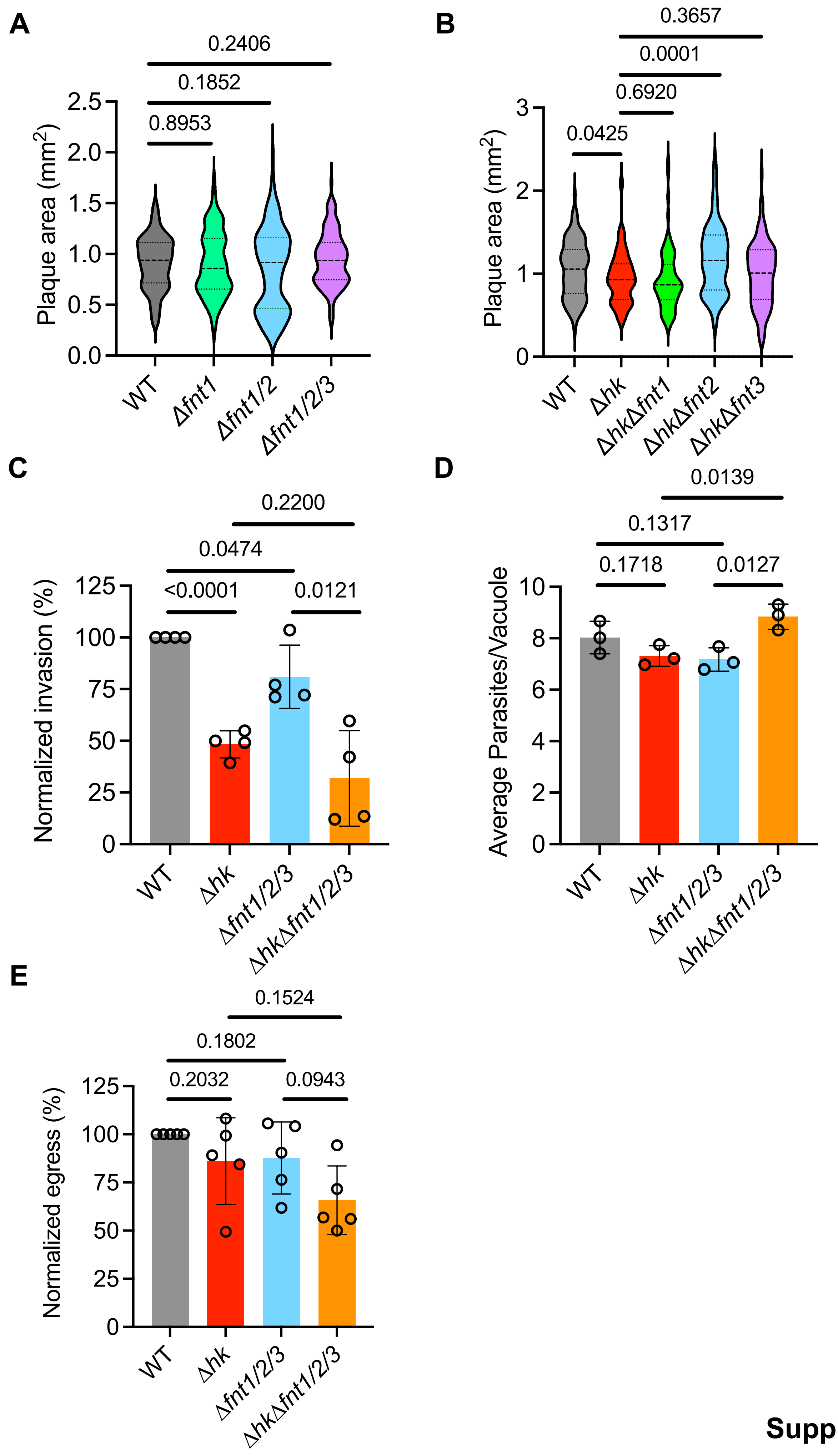


**Supplemental Figure 9**

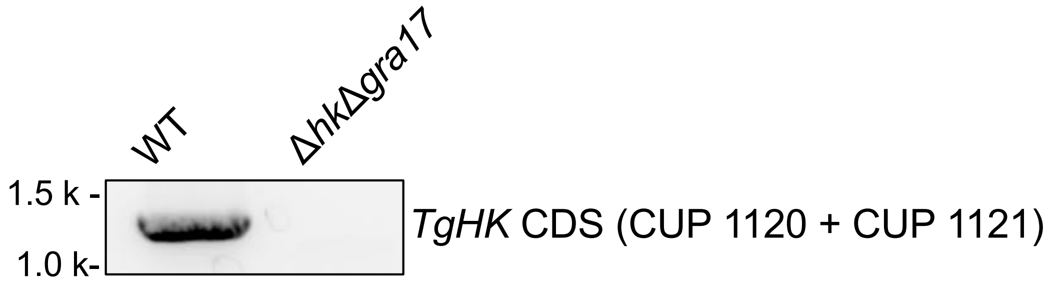


**Supplemental Figure 10**

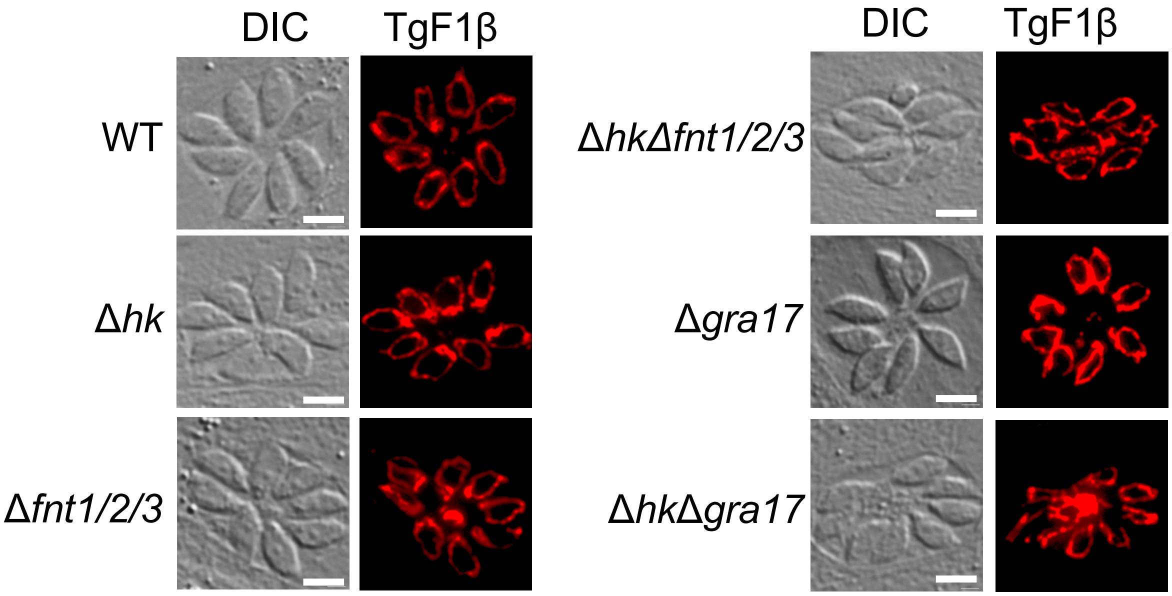


**Supplemental Figure 11**

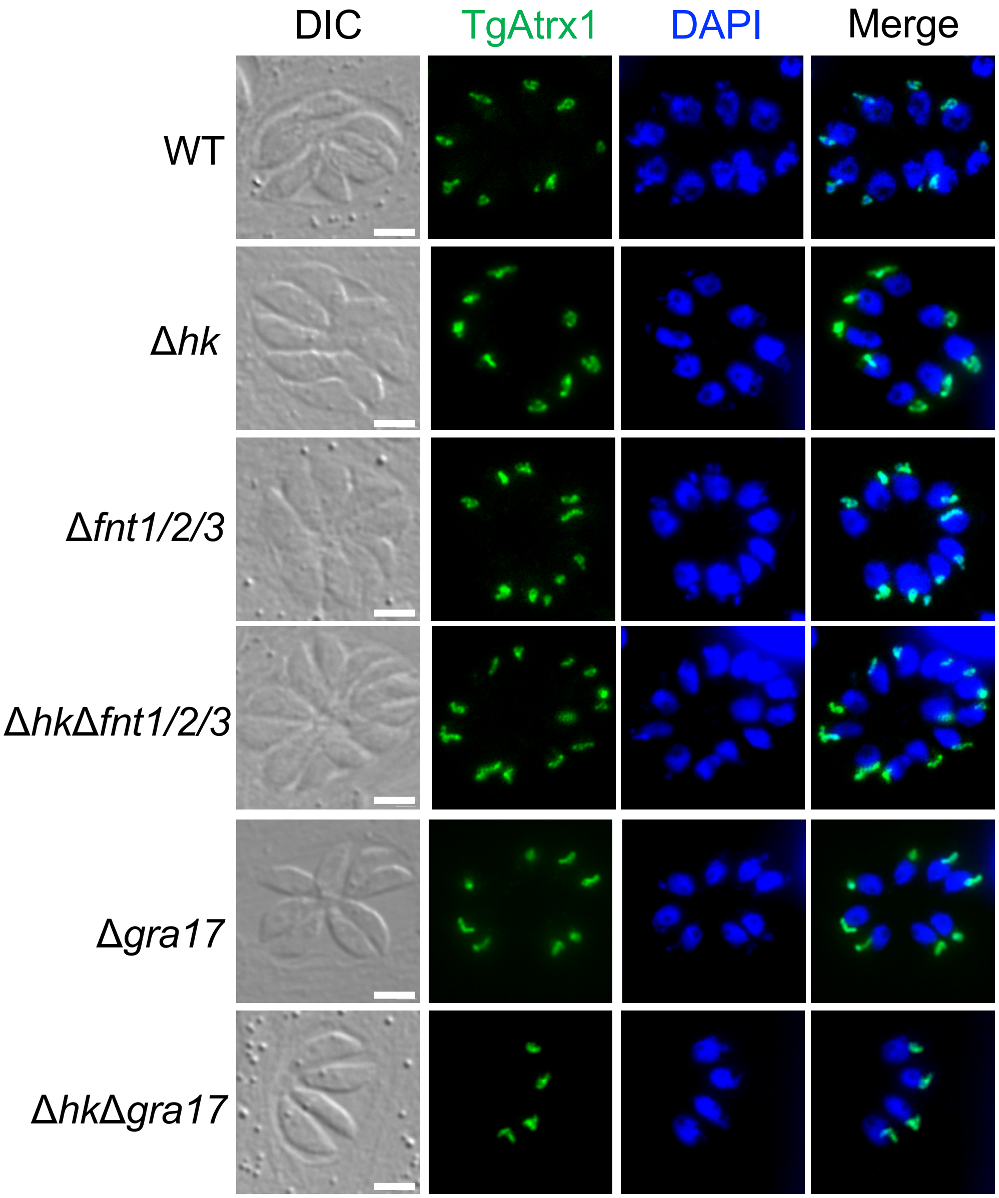


**Supplemental Figure 12**

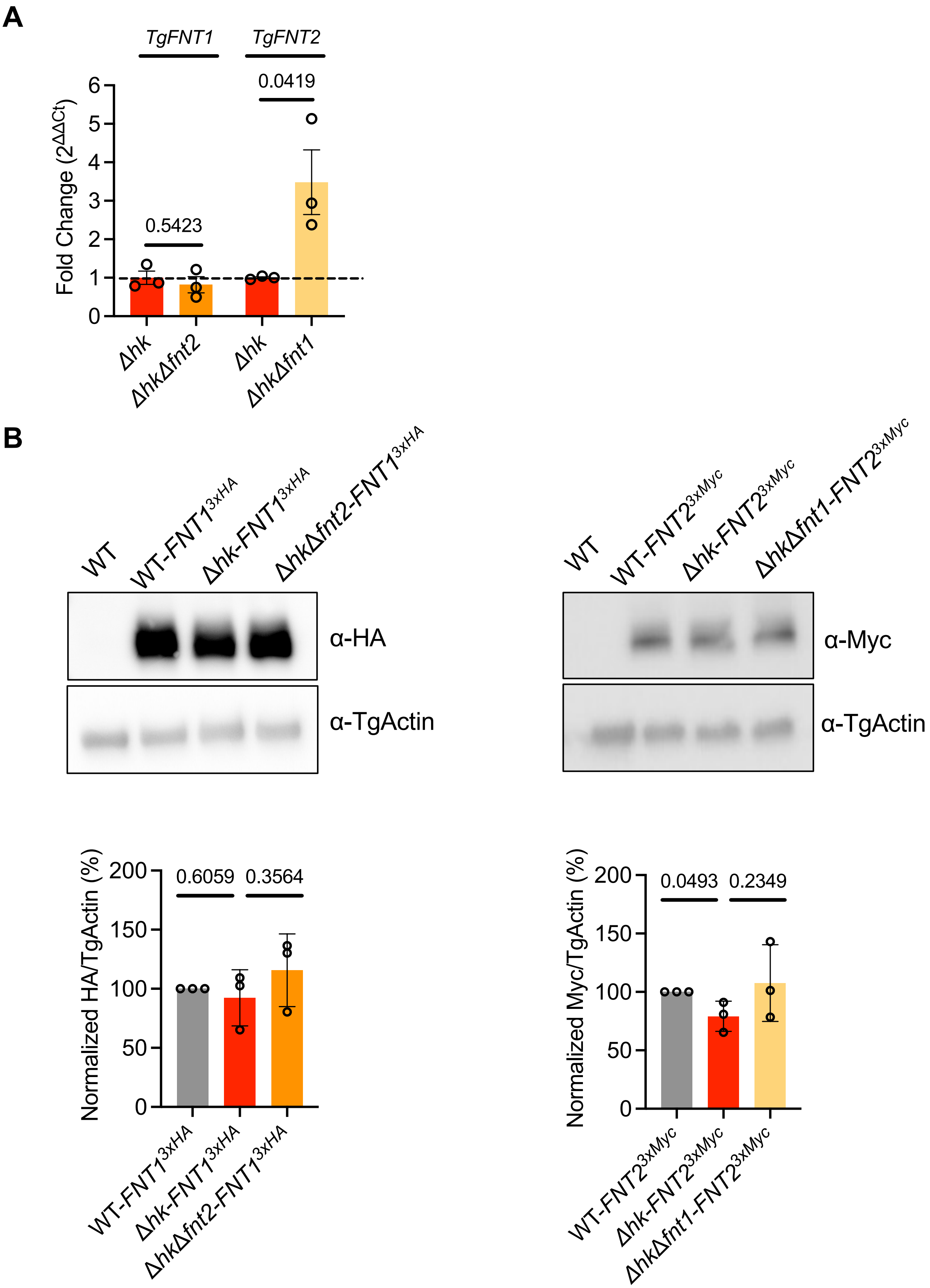


**Supplemental Figure 13**

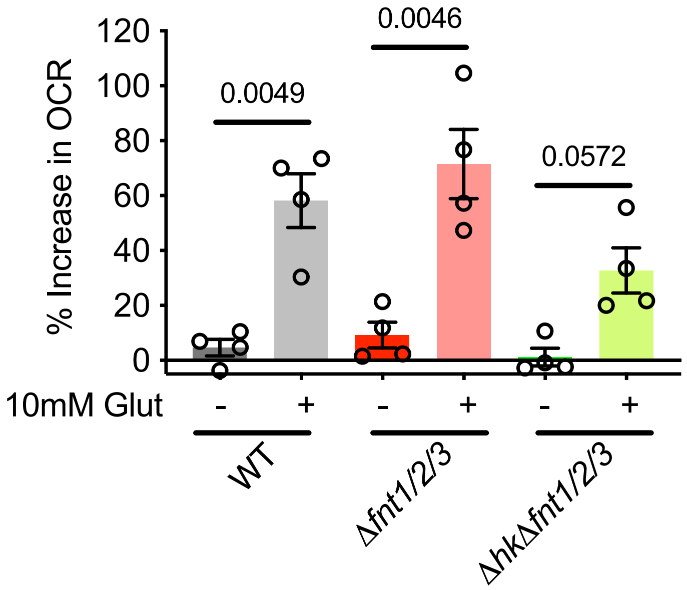


**Supplemental Figure 14**
